# Atoh7-independent specification of retinal ganglion cell identity

**DOI:** 10.1101/2020.05.27.116954

**Authors:** Justin Brodie-Kommit, Brian S. Clark, Qing Shi, Fion Shiau, Dong Won Kim, Jennifer Langel, Catherine Sheely, Tiffany Schmidt, Tudor Badea, Thomas Glaser, Haiqing Zhao, Joshua Singer, Seth Blackshaw, Samer Hattar

## Abstract

Retinal ganglion cells (RGCs), which relay visual information from the eye to the brain, are the first cell type generated during retinal neurogenesis. Loss of function of the transcription factor *Atoh7*, which is expressed in multipotent early neurogenic retinal progenitor cells, leads to a selective and near complete loss of RGCs. *Atoh7* has thus been considered essential for conferring competence on progenitors to generate RGCs. However, when apoptosis is inhibited in *Atoh7-*deficient mice by loss of function of *Bax*, only a modest reduction in RGC number is observed. Single-cell RNA-Seq of *Atoh7;Bax*-deficient retinas shows that RGC differentiation is delayed, but that RGC precursors are grossly normal. *Atoh7;Bax*-deficient RGCs eventually mature, fire action potentials, and incorporate into retinal circuitry, but exhibit severe axonal guidance defects. This study reveals an essential role for *Atoh7* in RGC survival, and demonstrates *Atoh7-*independent mechanisms for RGC specification.

## Introduction

The retina has six major classes of neurons that develop from a common progenitor cell pool during overlapping temporal intervals. Retinal ganglion cells (RGCs), the only projection neurons from the retina to the brain, are the first retinal cell type to be generated. RGC development in both zebrafish, mice, and humans has been shown to require the basic helix-loop-helix transcription factor atonal homolog 7 - *Atoh7 (Math5)* (Brown et al., 2001; Ghiasvand et al., 2011; Jarman et al., 1994; Khan et al., 2011; Khor et al., 2011; Macgregor et al., 2010; Prasov et al., 2012; Wang et al., 2001). *Atoh7* is conserved across all vertebrate species, and distantly related to *atonal*, which specifies the earliest-born neurons in *Drosophila* retina (Brown et al., 2001; Jarman et al., 1994; Kanekar et al., 1997; Prasov et al., 2012). *Atoh7*-deficient mice and zebrafish lack upwards of 95% of RGCs (Brown et al., 2001; Brzezinski et al., 2012; Kay et al., 2001; Wang et al., 2001) and likewise lack any visible optic nerve or functional connections from the retina to the brain (Brzezinski et al., 2005; Wee et al., 2002). Human mutations in *ATOH7* or its cis-regulatory regions have been associated with optic nerve agenesis or hypoplasia (Khan et al., 2012; Macgregor et al., 2010) and increased susceptibility to glaucoma (Fan et al., 2011; Ramdas et al., 2011). *Atoh7*-deficiency also disrupts development of retinal vasculature in both mice and humans, likely as an indirect result of the loss of RGCs (Edwards et al., 2012).

In mice, *Atoh7* is expressed in neurogenic retinal progenitor cells (RPCs) between E12 and P0, corresponding to the interval in which RGCs are generated (Brown et al., 1998; Clark et al., 2019; Feng et al., 2010; Pacal and Bremner, 2014; Prasov and Glaser, 2012; Rapaport et al., 2004; Sidman, 1960; Wang et al., 2001; Young, 1985). Upon cell fate specification, *Atoh7* expression is rapidly down-regulated in mouse RGC precursors (Clark et al., 2019; Miesfeld et al., 2018a) although expression persists in immature human RGCs (Aparicio et al., 2017; Lu et al., 2020). Genetic fate mapping indicates that *Atoh7*-expressing RPCs also give rise to other early-born retinal cells, including cone photoreceptors, horizontal and amacrine cells, and that generation of these cell types is increased in *Atoh7*-deficient mice (Brown et al., 2001; Brzezinski et al., 2005; Hufnagel et al., 2013; Wang et al., 2001). Ectopic expression of *Atoh7* alone, however, typically is not sufficient to drive RGC specification (Gan et al., 1999; Hufnagel et al., 2013; Mao et al., 2013, 2008; Ohnuma et al., 2002; Pan et al., 2008; Pittman et al., 2008; Prasov and Glaser, 2012; Wu et al., 2015), although misexpression of Atoh7 in Crx-expressing photoreceptor precursors was sufficient to rescue development of a limited number of RGCs (Prasov and Glaser, 2012).

These findings have suggested that *Atoh7* acts in neurogenic RPCs to confer competence to generate RGCs (Brzezinski et al., 2012; Mu et al., 2005; Yang et al., 2003) potentially in combination with as yet unidentified factors. Recent experiments have shown that when *Pou4f2* and *Isl1* are misexpressed under the control of the endogenous *Atoh7* promoter, this is sufficient to fully rescue the defects in RGC development seen in *Atoh7* mutants (Gan et al., 1999; Pan et al., 2008; Wu et al., 2015). This implies that *Atoh7* may act permissively to enable expression of these two factors in early-stage RPCs.

Other data, however, suggest that *Atoh7* may not be essential for RGC specification. Previous studies indicate that immature RGCs are present in *Atoh7*-deficient mice in embryonic retina, although at reduced numbers relative to controls (Brown et al., 2001; Brzezinski et al., 2012). Genetic fate mapping studies further raise questions about the necessity of *Atoh7* for RGC specification. Analysis of *Atoh7-Cre* knock-in mice reveal that only 55% of all RGCs are generated from *Atoh7*-expressing RPCs (Brzezinski et al., 2012; Feng et al., 2010; Poggi et al., 2005). Although this outcome may reflect inefficient activation of Cre-dependent reporter constructs, it may also imply that a subset of RGCs are specified through an *Atoh7*-independent mechanism that requires trophic support from *Atoh7*-expressing RPCs or *Atoh7*-derived RGCs.

To distinguish the role of *Atoh7* in controlling RGC specification and survival, we prevented RGC death in *Atoh7*-deficient mice by simultaneously inactivating the proapoptotic gene *Bax* (Chen et al., 2013; Knudson et al., 1995). Strikingly, we observed only a 25.2±0.9% reduction in adult RGC numbers in *Atoh7*^*−/−*^;*Bax*^*−/−*^ retinas relative to *Bax*^*−/−*^ controls. While mutant RGCs showed severe defects in formation of axonal projections and retinal vasculature, we found that ‘rescued’ RGCs expressed both *Pou4f2* and *Isl1* in addition to other markers of terminal differentiation, fired action potentials in response to light, and formed functional synapses with retinal neurons. Single-cell RNA-Sequencing (scRNA-Seq) analysis of *Atoh7;Bax*-deficient retinas shows that although RGC differentiation is delayed relative to wildtype, *Atoh7*-deficient RGCs express both *Pou4f2* and *Isl1.* This study demonstrates that, while *Atoh7* is required for both terminal differentiation and survival of RGCs, it is not necessary for specification of the majority of RGCs.

## Experimental Procedures

### Mice

Animals were housed and treated in accordance with NIH and IACUC guidelines, and used protocols approved by the Johns Hopkins University Animal Care and Use Committee (Protocol numbers MO16A212). *Atoh7*^*Cre/Cre*^ mice are a knock-in line where Cre recombinase replaced the entire *Atoh7* gene and was a gift from Dr. Lin Gan (referred to as Atoh7^−/−^) (Yang et al., 2003) (RRID:MGI:3717726). The Bax^tm1Sjk/tm1Sjk^ (*Bax*^*−/−*^) mice containing a neomycin cassette that replaces critical exons 2-5,were purchased from The Jackson Laboratory (Knudson et al., 1995)(JAX:002994, RRID:IMSR_JAX:002994). The *Bax*^*−/−*^ mice are unpigmented since the *Bax* gene is linked to the *Tyrosinase* (Tyr) and *Pink-eyed dilution* (*p*) gene by 21cM and 5cM respectively. The conditional Bax^tm2Sjk/tm2Sjk^ (Bax^fl/fl^) mice containing a LoxP sites flanking exons 2-4, were purchased from The Jackson Laboratory (JAX:006329, RRID:IMSR_JAX:006329) (Takeuchi et al., 2005). *Atoh7*^*tTA/tTA*^;B&I-EE mice are a combination of two genetic strains. In the first strain (*Atoh7*^*tTA/tTA*^), the tetracycline-responsive artificial transcription factor tTA replaces the *Atoh7* gene. In the absence of tetracycline, the tTA activates the tetracycline responsive element which is driving the expression of *Brn3b* and *Isl1* in the second strain (B&I-EE). Therefore, in effect the Atoh7 promoter will drive the expression of *Brn3b* and *Isl1*. This mouse line has been previously reported to rescue all reported effects of *Atoh7* loss of function, and was a gift from Dr. Xiuqian Mu (Wu et al., 2015)(MGI:5749708 and MGI:5749713). The *Crx>Atoh7* mice, a transgene that expresses the full-length *Atoh7* coding sequence under the control of the *Crx* promoter, was previously published (Prasov and Glaser, 2012) (MGI:5433215). A tdTomato Cre recombinase reporter mouse*Rosa26*^*tdTomAi14*^ (JAX:007914, RRID:IMSR_JAX:007914) (Madisen et al., 2010) was used to label cells in a Cre recombinase-dependent manner. The *Chx10-Cre* mouse line is a transgenic line purchased from The Jackson Laboratory, originally developed by Constance Cepko’s laboratory (JAX:005105, RRID:IMSR_JAX:005105) (Rowan and Cepko, 2004), expresses Cre recombinase broadly in all retinal progenitor cells from E10-E15.5. The *Opn4*^*taulacZ*^ mice were used to trace the ipRGC projections to the brain (Hattar et al., 2002). Throughout the manuscript, controls are heterozygous for both *Atoh7* and *Bax* (*Atoh7*^+/−^;*Bax*^*+/−*^), whereas *Atoh7*^−/−^ mice were also are heterozygous for *Bax* (*Atoh7*^−/−^;*Bax*^*+/−*^).

### Statistics

All statistical tests, apart from analysis of the scRNA-seq data, were performed in Graphpad Prism 6 (RRID:SCR_002798). The statistical tests used are listed in figure captions.

### Immunohistochemistry

Adult retinas from P40-P200 mice were obtained from by enucleating whole eyes, fixing for 30 minutes in 4% paraformaldehyde (PFA) diluted in PBS, dissecting to remove the cornea and lens, dissecting the retina from the RPE, and antibody staining proceeded in a 24-Multiwell Cell Culture Plate (Corning #353047). Retinas were blocked in 500 μl of PBS containing 0.3% Triton X100 and 6% goat serum for 2 hours at room temperature. Several antibodies were used in this study (dilutions are between brackets): Mouse IgG1 anti-Brn3a (Millipore Cat# MAB1585 RRID:AB_94166) (1:250), Rabbit anti-RBPMS (GeneTex Cat# GTX118619 RRID:AB_10720427) (1:250), Rabbit anti Brn3b (Badea et al., 2009) (1:100), Mouse anti-Smi32 (non-phosphorylated anti-Neurofilament H (NF-H)) (BioLegend Cat# 801701 RRID:AB_2564642) (1:500), Mouse anti Neurofilament Medium (Thermo Fisher Scientific Cat# 13-0700, RRID:AB_2532998) (1:500), Chicken anti Neurofilament Heavy (Millipore Cat# AB5539, RRID:AB_11212161) (1:250), Mouse anti-GFAP (Sigma-Aldrich Cat# C9205 RRID:AB_476889) (1:1000), GS-IB4 (Molecular Probes Cat# I21411 also I21411 RRID:AB_2314662) (1:250), Rabbit anti-Pax2 (BioLegend Cat# 901001 RRID:AB_2565001) (1:100), Mouse anti-Tuj1 (R and D Systems Cat# MAB1195 RRID:AB_357520) (1:200), Rabbit anti DsRed Takara Bio (Cat# 632496, RRID:AB_10013483) (1:250), Mouse anti-Islet1 (DSHB Cat# 40.2D6 RRID:AB_528315) (1:200), and Rabbit anti-Opn4 (Advanced Targeting Systems Cat# UF006, RRID:AB_2314781) (1:500). The appropriate antibodies were diluted in blocking solution and incubated for two days at 4°C. Retinas were then washed in three changes of PBS, 15 minutes each, then placed in the appropriate Alexa Fluor secondary antibody (Invitrogen) (1:500) overnight at 4°C. Retinas were washed in 200 μl of PBS containing 1xDAPI, then washed three times in PBS for 15 minutes each, and mounted flat on slides in VectaShield (Vector Labs, RRID:AB_2336789).Regionalized dissections were done as follows; before enucleation, the most nasal part of the sclera was marked with a cauterizer. This mark was used during the dissection to make a marking incision into the retina, following the above staining protocol. Retinas were imaged on a Zeiss LSM 700 or 800 Confocal at the Johns Hopkins University Integrated Imaging Center Core Facility (RRID:SCR_016187).

For embryonic studies, developing embryos harvested at E12.5 and E14.5 were washed in a Petri dish with sterile PBS three times for 10 minutes. Tail was used for genotyping. The heads were fixed in 4% PFA for 30 minutes and then cryoprotected in 30% sucrose at 4°C overnight, frozen in OCT, and sectioned at 18 μm thickness using a cryostat. Sections were dried at 30°C for 15 minutes and then washed 10 minutes in three changes of PBS. Sections were then blocked and stained as above in a humidified chamber overnight. Sections were then mounted and imaged as described.

### Cell density analysis

All cell counting was done manually. To confirm the reproducibility of the cell counts, randomly selected selections from each sample were counted twice, and counts were consistently found to be essentially identical. Density was calculated as the number of cells per area. All measurements and cell number analysis was done manually in ImageJ (Fiji, RRID:SCR_002285) and Adobe Photoshop CS6 (RRID:SCR_014199).

In adult flat-mounted retinas, density of RGCs was calculated by obtaining at least 4 representative images at 40x of 600 μm × 600 μm with 1 μm optical sections. Optical sections were projected together with maximum intensity, including cells only in the retinal layers of interest. Representative images were taken similarly across all genotypes without *a priori* knowledge of the genotype. However, some genotypes contain marked phenotypic differences, which include a pigment mutation linked to the *Bax* locus, drastic RGC number reduction as in the *Atoh7*^*−/−*^, and/or misguided axons. Three representative areas of each retina were averaged for the density analysis

In E12.5 embryonic retinal sections, a representative confocal image was taken at 40x of 600 μm × 600 μm with optical sections of 1 μm projected together with maximum intensity. The sections chosen for analysis were all positive for Pax2+ optic nerve head cells, as the central retina contains the earliest-born RGCs. At least two sections with matching criteria were analyzed for each E12.5 embryo. Density was calculated by dividing the number of Brn3a+ RGCs and dividing by the area of the retina. To limit the analysis to the RGC neurogenic zone, we limited the quantification to the leading edge of RGC genesis. The percentage of mature Brn3a+ RGCs at E12.5 were determined by counting their number within the ganglion cell layer (GCL). The number of mature Brn3a+ RGCs was then divided by the number of total Brn3a+ RGCs in a section, and then averaged across all sections. This ratio represents the number of Brn3a+ RGCs already in the nascent ganglion cell layer versus RGCs migrating through the neuroblast layer to the ganglion cell layer.

### Multielectrode array recordings

Mice were dark-adapted for 1-2 hours before being sacrificed and dissected under dim red light. Retinas were isolated in Ames’ medium (Sigma) bubbled with 95% O2/5% CO2 (carbogen) at room temperature, trimmed into small rectangles, and then placed on a 6×10 perforated multielectrode array (Multichannel Systems, Tübingen, Germany), ganglion-cell-side down. Tissue was perfused with Ames’ bubbled with carbogen and kept at 32°C throughout the experiment. Data acquisition was performed using the MC_Rack software (ALA Scientific Instruments, Inc.), at a 50-kHz sampling rate. An offline spike sorter (Plexon Inc) was used for spike sorting.

UV stimuli (I_mean_ ≈ 5 × 103 photons/cm2/s, 398 nm) were generated through a modified DLP projector (HP Notebook Projection Companion Projector, Model: HSTNN-FP01) (frame rate = 60 Hz) and were delivered through an inverted microscope objective. All stimuli were programmed using the *Psychophysics Toolbox* in Matlab (The Mathworks, Natick, MA). Stimuli include: (1) 120-s, 1-Hz full-field square-wave flash (100% Michelson contrast); (2) 10-min Gaussian white noise (GWN) flickering checkerboard (pixel size = 44.77 μm); (3) 10-min spatially correlated “cloud” stimulus that was generated by low-pass filtering the GWN. The cloud stimulus introduced dark and bright areas of a range of scales within each frame, with the purpose of driving large spatial receptive fields.

Analysis was first performed using custom-written Matlab (MATLAB R2014b) codes, the results later were exported and edited in Adobe Illustrator CS6. For each cell, the peristimulus time histogram (PSTH) of responses to square-wave flash was calculated using 10-ms bins. Spatial and temporal receptive fields were identified based on noise data using a nonlinear model previously described in detail (McFarland et al., 2013; Shi et al., 2019).

Fewer cells were recorded from *Atoh7*^*Cre/Cre*^*;Bax*^*−/−*^ mice compared to the wildtype and Bax^−/−^, as the nerve fiber layer (NFL) and retinal vasculature are improperly developed and thus provide an insulating layer that needs to be removed in order to obtain high quality recordings. No cells were recorded from *Atoh7*^*Cre/Cre*^ mice, due to the >99% reduction in RGC numbers.

### Pupillary light response (PLR)

PLR experiments were performed on mice that were dark adapted for at least 1 hour prior to any experiment. PLR was measured by gently restraining the mice by hand (without anesthesia) and exposing them to a cool white light LED bulb (6500K, light intensity: 15 W/m^2^, MR16, SuperBrightLEDs.com) that was directed at one eye using a gooseneck arm of a dissecting microscope light source. Constriction of the pupil was recorded using a Sony Handycam camcorder (FDRAX33) from either the contralateral or ipsilateral eye to the light source. The baseline pupil size of each mouse was first recorded for at least 5 seconds using an infrared light source, following which the white LED bulb was turned on for at least 30 seconds. Video recordings were analyzed by creating screen shot images in Joint Photographic Experts Group format (jpg) of the pupil prior to and during light stimulation using VLC media player (https://www.videolan.org/vlc/). Pupil area was then quantified in ImageJ (Fiji, RRID:SCR_002285). To determine the relative pupil area, pupil size during the light stimulation was divided by pupil size prior to light stimulation.

### Tissue dissociation for generation of single-cell suspensions

Eyes were enucleated from E14 time-pregnant animals and placed directly into ice-cold 1X PBS. Retinas were dissected in cold 1X PBS, and then placed into 200 μl of cold HBSS per 2-3 retinas. Tissue dissociation was induced through addition of an equivalent volume of Papain solution (1 ml - 700 μl reagent grade water, 100 *μ*l fresh 50 mM L-Cysteine (Sigma), 100 μl 10 mM EDTA, 10 μl 60 mM 2-mercaptoethanol (Sigma), with Papain added to 1 mg/ml (1:10 dilution of 10 mg/ml Papain solution; Worthington)). Papain-retina mixture was placed at 37°C for 10 minutes with slight trituration every 2-3 minutes. Enzymatic dissociation was halted through addition of 600 μl of Neurobasal Media + 10% FBS for every 400ul of dissociation solution. DNA from lysed cells was removed using 5 μl RNAse-free DNAse I (Roche) for every 1 ml dissociate and incubated at 37°C for 5 minutes, followed by slight trituration using a 1 ml pipette. Cells were pelleted after centrifugation (300 RCF for 5 minutes at 4°C), followed by resuspension in 2-3 ml of Neurobasal media supplemented with 1% FBS. The final solution was passed through a 50 μm filter to remove cellular aggregates and undissociated debris.

### 10x Genomics Sequencing and Analysis

Single cell RNA sequencing of dissociated retinal cells from E14 *Atoh7*^*-/+*^*;Bax*^*−/−*^ and *Atoh7*^*−/−*^*;Bax*^*−/−*^ was performed using 10x Genomics Chromium 3’ v2 platform (PN-120223) (Pleasanton, CA), followed by sequencing using the NextSeq500 platform with default 10x sequencing parameters (R1 - 26bp; R2 - 98bp; i7 - 8bp). Single-cell analysis of the wildtype (WT) E14 developing mouse retina was obtained from previously reported samples obtained from GEO (GSE118614); data obtained using similar isolation protocols are described previously (Clark et al., 2019).

### Gene Set Usage Pattern Discovery with scCoGAPS

CoGAPS v.3.5.6 (Sherman et al., 2019; Stein-O’Brien et al., 2019) was used to find patterns of gene set usage by Neurogenic and Retinal Ganglion cells. The expression matrix used as input was normalized to 10,000 counts per cell, subsetted down to 5235 most highly variable genes and log2 transformed. The CoGAPS parameters used are: singleCell=TRUE, nPatterns = 30, nIterations = 50000, distributed = single-cell, sparseOptimization = True, seed = 803L, and nSets = 10. The final number of patterns stabilized at 31.

### Identification of *Atoh7*-Dependent Genes

Data was subsetted down to neurogenic RPCs and RGCs. Monocle’s differential gene test was conducted between control (WT and *Bax*^*−/−*^) and Atoh7 mutants (Atoh7^−/−^ Atoh7^−/−^;Bax^−/−^) as such:

differentialGeneTest(dat[genes expressed in >=10 cells], fullModelFormulaStr = ‘~(Atoh7 genotype) + Total_mRNAs’, reducedModelFormulaStr = ‘~Total_mRNAs’,cores=4).

### Pseudotime Analysis between Genotypes

Scanpy v1.4 (Wolf et al., 2018) was first used to assign diffusion pseudotime values (Haghverdi et al., 2016) to cells in the retinal ganglion cell trajectory. Cell types included in this final dataset were restricted to retinal ganglion cells, primary RPCs and neurogenic RPCs. To preprocess the dataset, genes <10 counts were removed, and the expression matrix was normalized to 10,000 counts/cell and log-transformed. Highly variable genes used for ordering were identified using Scanpy’s ‘highly_variable_genes’ function with default parameters except flavor=’cell_ranger’ and n_top_genes=3000. 50 principal components were calculated using default PCA parameters with random_state=123456. To compute the neighborhood graph with the batch effect of genotype removed, we used BBKNN with batch_key = “Genotype” and neighbors_within batch=3 (Polański et al., 2019). 10 diffusion components were then computed and used for input to assign diffusion pseudotime values with an RPC cell as root. To find genes differentially expressed between the developmental trajectories of the WT and *Atoh7*^*Cre/Cre*^*;Bax*^*−/−*^ genotypes, Monocle’s differential gene test (Qiu et al., 2017; Trapnell et al., 2014) was performed in R, on neurogenic RPCs and RGCs of WT and *Atoh7*^*Cre/Cre*^*;Bax*^*−/−*^ null genotypes:

differentialGeneTest(dat[genes expressed in >=10 cells], fullModelFormulaStr = ‘~sm.ns(Pseudotime,df=3)*Genotype+Total_mRNAs’, reducedModelFormulaStr = ‘~sm.ns(Pseudotime,df=3)+Genotype+Total_mRNAs’,cores=3)

### Data Availability

Processed (expression, gene (featureData), and cell (phenoData) matrices) and raw sequence information (.bam files) are available for direct download through GEO GSE148814.

### *In situ* Hybridization

Developing embryos harvested at E14.5 were washed in petri dishes filled with sterile DEPC-treated PBS at least three times. The head of the embryo was plunged into OCT and then immediately frozen and stored at −80°C until needed, and the tail was used for genotyping. 20 μm sections were taken using a cryostat. Sections were allowed to dry to slides for a few hours and then were stored at −80°C until needed. *In situ* hybridization was performed as previously described (Shimogori et al., 2010).

## Results

### *Atoh7* promotes RGC survival, but RGC specification is largely *Atoh7*-independent

In the absence of *Atoh7,* there is an increase in apoptosis of both *Atoh7*-derived cells across embryonic retinal development and non-*Atoh7-*derived cells in the GCL at E16.5 and E17.5 (Feng et al., 2010; Prasov and Glaser, 2012). These data suggest that *Atoh7* may promote RGC survival in both a cell-autonomous and cell non-autonomous manner. To better understand the role *Atoh7* plays in RGC development, independent of its role in RGC survival, we disrupted both *Atoh7* and the proapoptotic *Bax* gene, in order to inhibit apoptosis in the retina.

We used *Atoh7*^*Cre/Cre*^ mice, in which the *Atoh7* coding sequence is replaced with Cre recombinase via targeted recombination, generating a null allele, to analyze Atoh7 function (Yang et al., 2003). We first examined the expression of RBPMS and Isl1, both of which are broadly expressed in RGCs, in *Atoh7*^*Cre/Cre*^*;Bax*^*−/−*^ mice (Figure 1A-C) (hereafter referred to as *Atoh7*^*−/−*^*;Bax*^*−/−*^ mice). *Isl1*, a LIM family homeodomain transcription factor, is necessary for RGC development and maintenance in adulthood (Mu et al., 2008; Pan et al., 2008), and is expressed in mature RGCs, bipolar, and amacrine cells. Using anti-Rbpms to selectively label all RGCs, we observed an 159±7% (Figure 1A,B) increase in RGC number in *Bax*^*−/−*^ retinas, in line with previous results indicating that RGCs undergo extensive levels of apoptosis during development. However, we observe only a 25.2±0.9% reduction in RGCs in *Atoh7*^*−/−*^*;Bax*^*−/−*^ relative to *Bax*^*−/−*^ retinas. This contrasts with the 99.54±0.12% reduction in RGCs in the *Atoh7*^*−/−*^ compared to controls (Figure 1A,B). Similar results were observed for anti-Isl1 cells in the GCL (Figure 1A,C).

**Figure 1.**
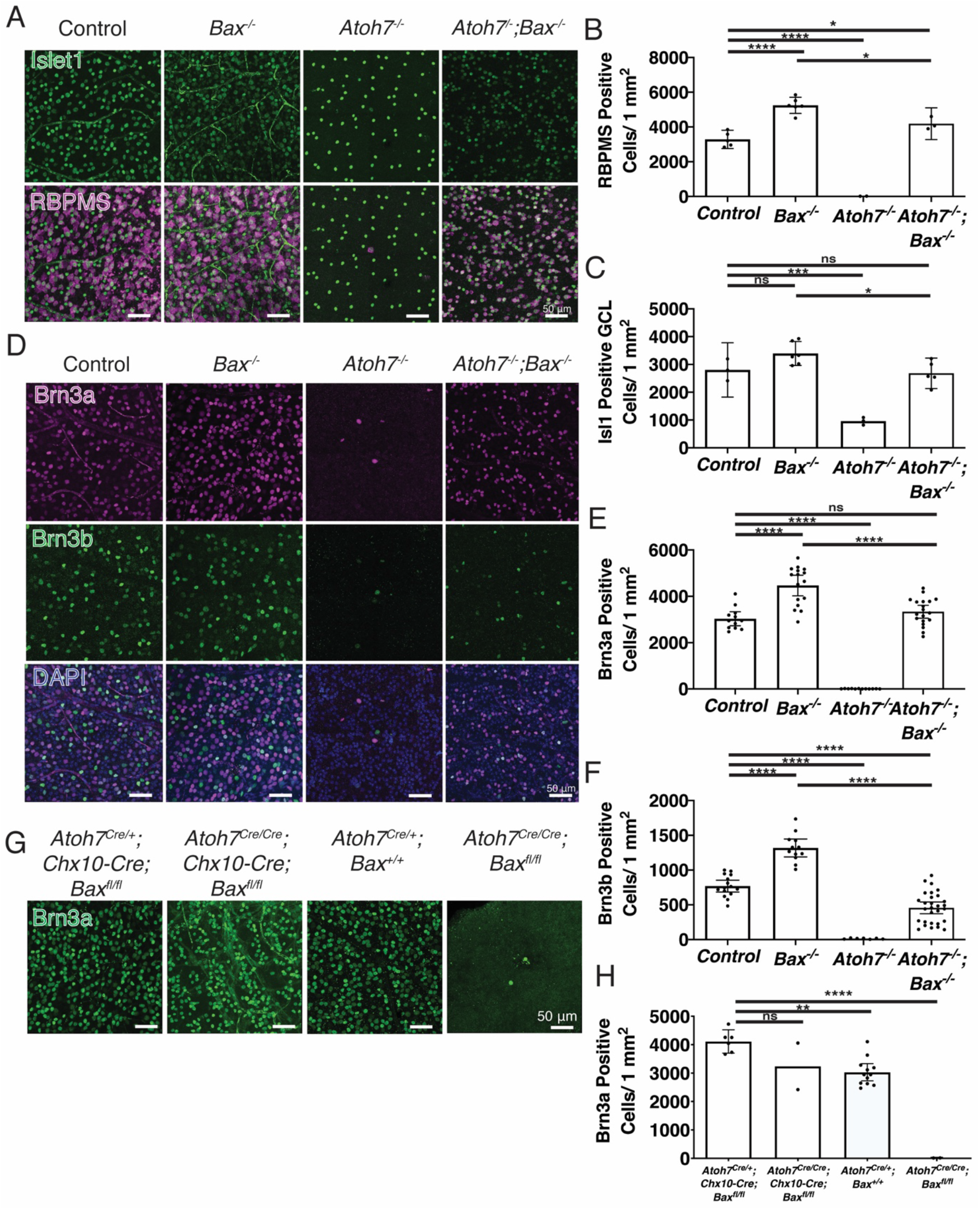
*Atoh7*-independent development of RGCs. (A-C) We observed a 25.2±0.9% and 21±3% reduction in RBPMS+ RGC density or Isl1+ ganglion cell layer (GCL) cells when comparing *Atoh7*^*−/−*^*;Bax*^*−/−*^ to *Bax*^*−/−*^ mice. (D,E,F,G,H) Brn3a and Brn3b positive RGC density are only moderately reduced when apoptosis is blocked in *Atoh7*^*−/−*^*;Bax*^*−/−*^ mice. (G,H) Brn3a positive RGC numbers are rescued when apoptosis is blocked in all neural retinal progenitor cells, when *Bax*^*lox/lox*^ is crossed to the *Chx10-Cre* transgene, which is expressed in all RPCs. However, when *Bax* is specifically removed in *Atoh7-Cre* knock-in mice, Brn3a RGCs are not rescued. Mean ± 95% confidence intervals. Statistical significance tested by one-way ANOVA with Tukey’s post test for multiple comparisons **** p < 0.0001.

Specific RGC markers, Brn3a (Pou4f1) and Brn3b (Pou4f2), were used to quantify RGCs. For wildtype and *Bax*^*−/−*^lines, Brn3a and Brn3b numbers were similar to published reports (Figure 1D-F) (Prasov and Glaser, 2012; Rodriguez et al., 2014; Wang et al., 2001; Xiang et al., 1995). In the *Atoh7*^*−/−*^ line a 99.72±0.17% reduction in Brn3a RGC density was observed. However, in *Atoh7*^*−/−*^*;Bax*^*−/−*^ mice, the Brn3a RGCs were substantially rescued in the *Atoh7*^*−/−*^ background and remained into adulthood. 74.7±0.9% of Brn3a RGCs are rescued in *Atoh7*^*−/−*^*;Bax*^*−/−*^ mice relative to *Bax*^*−/−*^ levels in adult, and RGCs display normal distribution across the entire retina (Figure 1D,E, Supplemental Figure 1A,A’). Interestingly, Brn3b RGCs were also increased in *Atoh7*^*−/−*^*;Bax*^*−/−*^ relative to *Bax*^*−/−*^ retinas, but to a lesser extent than the Brn3a (28.8±5.9%; Figure 1D,F, Supplemental Figure 1B,B’). Expression of *Isl1*, *Brn3b*, and, to a lesser extent *Brn3a* have previously reported to require *Atoh7* (Mu et al., 2008, 2004, 2005; Pan et al., 2008; Rodriguez et al., 2014; Wu et al., 2015). However, our data indicates that the expression of *Isl1*, *Brn3a,* and *Brn3b* expression in RGCs can occur independent of *Atoh7* (Figure 1).

To investigate the extent to which rescued RGCs resembled wildtype neurons, we examined the expression of markers of major classes of mature RGCs. We investigated the prevalence of intrinsically photosensitive RGCs (ipRGCs) within *Atoh7*^*−/−*^*;Bax*^*−/−*^ retinas. During the development of ipRGCs, a majority of cells express Brn3b, although in some cases only transiently (Chen and Hattar, 2012). To determine the percentage of rescued ipRGCs in *Atoh7*^*−/−*^*;Bax*^*−/−*^ mice, we used a melanopsin antibody that predominantly labels the high melanopsin-expressing M1 and M2 ipRGC populations. We observe that 34.1% of ipRGCs are rescued in *Atoh7*^*−/−*^*;Bax*^*−/−*^ mice relative to *Bax*^*−/−*^, proportions similar to the fraction of Brn3b-positive RGCs in wildtype (Figure 1D,F, Supplemental Figure 1C-F). This indicates that Brn3b-positive ipRGCs differentiate normally in the absence of *Atoh7*.

To eliminate the possibility that global loss of function of *Bax* caused a non-specific rescue of RGC development, we tested the effects of retina-specific conditional mutants of *Bax*. Using *Chx10-Cre;Atoh7*^*Cre/Cre*^*;Bax*^*fl/fl*^ (Figure 1G,H). In this model, *Bax* is selectively disrupted in RPCs beginning at E10-10.5 (Rowan and Cepko, 2004). Removal of *Bax* from all RPCs shows RGC development to the same extent as in *Atoh7*^*Cre/Cre*^*;Bax*^*−/−*^(Figure 1G, H). In contrast, *Atoh7*^*Cre/Cre*^*;Bax*^*fl/fl*^ mice did not show any detectable rescue of RGC development (Figure 1G,H). This suggests that the survival-promoting effects of *Atoh7* may take place only during a narrow temporal window, and that Cre-mediated deletion of *Bax* in the *Atoh7*^*Cre/Cre*^ line may not occur rapidly or efficiently enough to rescue RGC development.

In both wildtype and *Atoh7*^*−/−*^*;Bax*^*−/−*^ animals, 34±11% and 34±26% of RGCs are derived from Atoh7-expressing cells, a finding which independently confirms similar lineage tracings in previous studies (Supplemental Figure 2A-C) (Brzezinski et al., 2012; Feng et al., 2010). This indicates that while RGCs that are normally derived from *Atoh7*-expressing neurogenic RPCs are reduced in the absence of *Atoh7*, *Atoh7* is not required for their specification. Furthermore, RGCs that are derived from non-*Atoh7-* expresing RPCs require *Atoh7* for survival through a non-cell autonomous mechanism.

### RGCs specified in the absence of *Atoh7* generate light-driven photic responses transduced from the outer retina

As we determined that loss of Bax in the *Atoh7*^*−/−*^ background rescues RGC specification, we next sought to investigate the degree to which RGCs differentiated in the absence of *Atoh7*. One hallmark of RGC differentiation is formation of presynaptic contacts and the generation of light-induced action potentials. We used multi-electrode array recordings to test the light responsiveness of the RGCs within the various mutant models. Spatiotemporal noise stimuli were used to activate the retina with a mean excitation of 398 nm (*I*_mean_ ≈ 5 × 10^3^ photons/cm^2^/s), a wavelength that predominantly activates S-cones (Wang et al., 2011), and does not activate Opn4-expressing ipRGCs (Berson et al., 2002; Do, 2019; Do and Yau, 2010; Hattar et al., 2002). We observed an identical stimulus-response profile in wildtype and *Bax*^−/−^ RGCs (Figure 2A): an expected result given that *Bax*^−/−^ mice display normal visual responses within the Morris water maze test (Chen et al., 2013). The population of RGCs included cells with similar light-evoked responses as ON sustained, ON transient, ON/OFF transient, OFF transient, or OFF sustained RGCs(Figure 2A, Supplemental Figure 3). The spatial receptive fields of the *Atoh7*^*−/−*^*;Bax*^*−/−*^ RGCs were also slightly smaller than normal (Atoh7^−/−^;Bax^−/−^ average 191±37.1 μm [n=56 from 8 mice], control average 212±42.3 μm [n=92 from 5 mice] and Bax^−/−^ average 223±42.4 μm [n=96 from 4 mice]) (Figure 2B) and the kinetics of the light-driven responses in the *Atoh7*^*−/−*^*;Bax*^*−/−*^ retinas slightly slower than those of control and Bax^−/−^ cells (Figure 2C).

**Figure 2.**
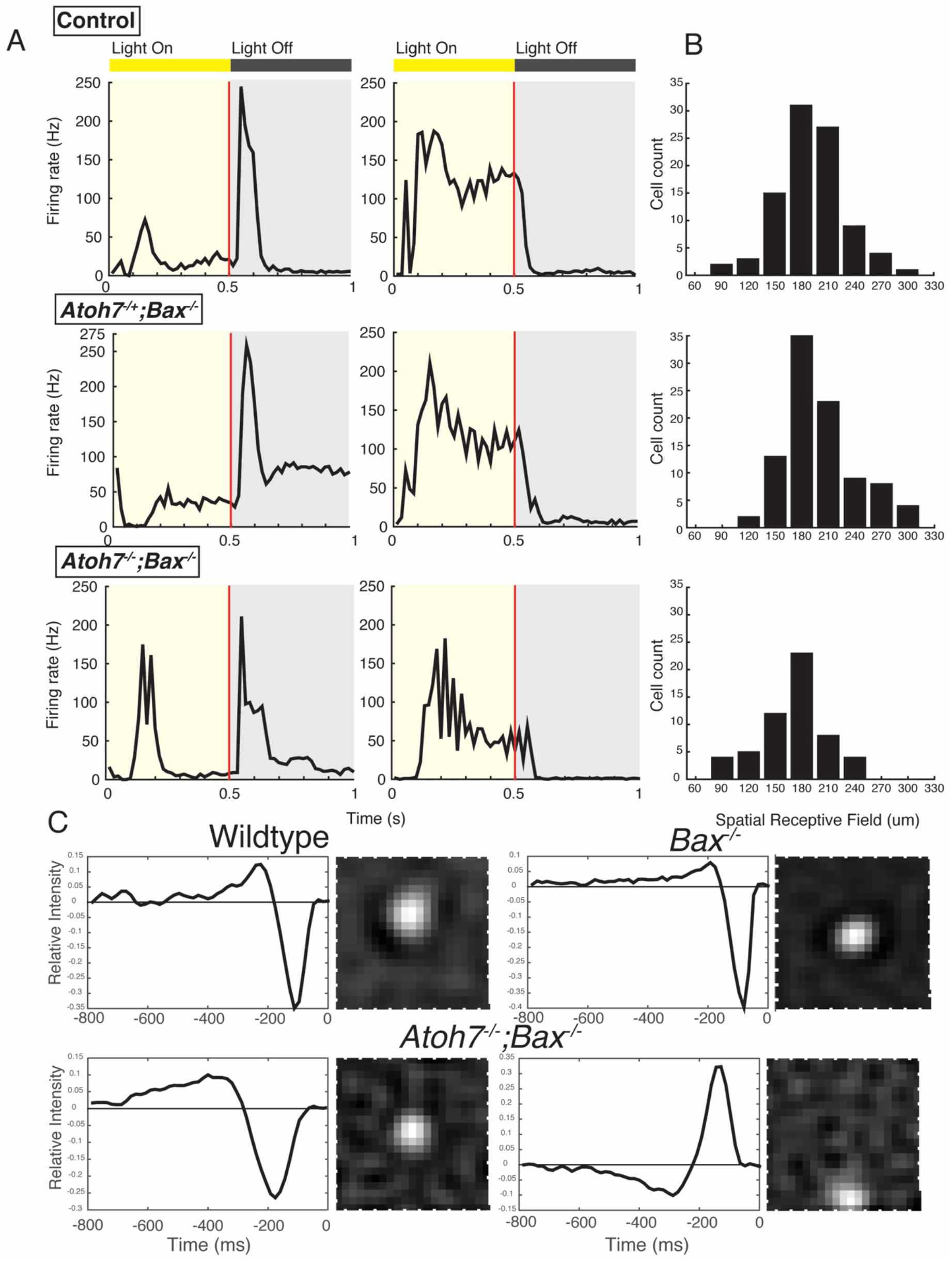
*Atoh7* is not required for normal retinal wiring and electrophysiological function. Cells from *Atoh7*^*+/−*^*;Bax*^*−/−*^, and *Atoh7*^*−/−*^*;Bax*^*−/−*^ mice responded to light similarly as those from wild type. (A) Two different examples, corresponding to two different sets of RGCs, per genotype of peristimulus time histogram averaged from 120 repetition of 1-Hz square wave flash: WT (upper panel), *Atoh7*^*+/−*^*;Bax*^*−/−*^ (middle), and Atoh7^−/−^;Bax^−/−^ (lower panel). (B) Distribution of the spatial receptive field measured using white noise flickering checkerboard: WT (upper, cell count = 92), *Atoh7*^*+/−*^*;Bax*^*−/−*^ (middle, cell count = 94), and *Atoh7*^*−/−*^*;Bax*^*−/−*^ (lower, cell count = 56). (C) the peristimulus time histogram (PSTH) of responses to square-wave flash was calculated using 10-ms bins. Mice assayed at P30. one-way ANOVA, followed by Dunnett’s test, *p* < 0.05, between *Atoh7*^*−/−*^*;Bax*^*−/−*^ and wildtype.

The linear analysis used for the MEA data can only reveal an averaged STA, which could be a compression of multiple receptive fields. Thus those having multiple receptive fields such as ON-OFF cells could potentially be hidden with this method and require more sophisticated analysis strategy (Shi et al., 2019). However, diversity in PSTH profiles is clearly observed, suggesting that different cell types coexist in *Atoh7*^*−/−*^*;Bax*^*−/−*^ mice. We chose not to perform more complex analysis such as classifying each cell into known cell types due to the relatively small sample size. These will be intriguing questions to address when larger amounts of data are made accessible. The near-normal properties of the *Atoh7*^*−/−*^*;Bax*^*−/−*^ RGCs show that RGCs present in the *Atoh7*^*−/−*^*;Bax*^*−/−*^ are wired properly to the outer retina, specifically the S-cones, and receive normal circuit input.

### RGC axon guidance is *Atoh7*-dependent

While RGCs in *Atoh7*^*−/−*^*;Bax*^*−/−*^ animals appropriately respond to detection of visual stimuli by the outer retina, the ability of these RGCs to form postsynaptic connections in the brain is compromised. We observed a substantially reduced pupillary light response in *Atoh7*^*−/−*^*;Bax*^*−/−*^ animals compared to controls (Figure 3B,C). To determine the cause of behavioral deficits, we assessed the distribution of RGC axons. Immunostaining for Smi32 (non-phosphorylated Nfh) (Figure 3A,D, Supplemental Figure 4C), Nfh, and Nfm (Supplemental Figure 4A,B) was used to evaluate RGC axonal integrity, and showed normal architecture in wildtype and *Bax*^*−/−*^ mice. We were surprised to find that the <1% of RGCs that survive in the *Atoh7*^*−/−*^ showed severe guidance defects, in that they fasciculate, come in close proximity to where the optic disc should be, seem to overshoot the optic disc, and then continue to extend within the retina. The great majority of *Atoh7*^*−/−*^ RGC axons fail to correctly target the optic disc, with only a few axons exiting and forming a rudimentary optic nerve, leading to a severely reduced PLR (Figure 3B,C, Supplemental Figure 5A-C). The gross misguidance of axons was also observed in *Atoh7*^*−/−*^*;Bax*^*−/−*^ mice (Figure 3A,D). As in *Atoh7*^*−/−*^ mice, RGC axons in *Atoh7*^*−/−*^*;Bax*^*−/−*^ mice fasciculate, fail to correctly target the optic disk, and form tracts that extend around the retina, with only rudimentary optic nerves observed. This guidance defect did not result from defective formation of the optic disc, as Pax2-positive glial cells that mark this structure are present in the *Atoh7*^*−/−*^*;Bax*^*−/−*^ at E12.5 (Supplemental Figure 6A,B) and E14.5 (data not shown). This demonstrates that the optic disk is present, but the axons are unable to find their way out of the retina in large numbers. These findings are reminiscent of previous reports in zebrafish, in which morpholino-mediated disruption of *atoh7* expression in early-stage RPCs disrupted the correct targeting of axons of later-born, *atoh7*-positive RGCs to the optic tectum (Pittman et al., 2008).

**Figure 3.**
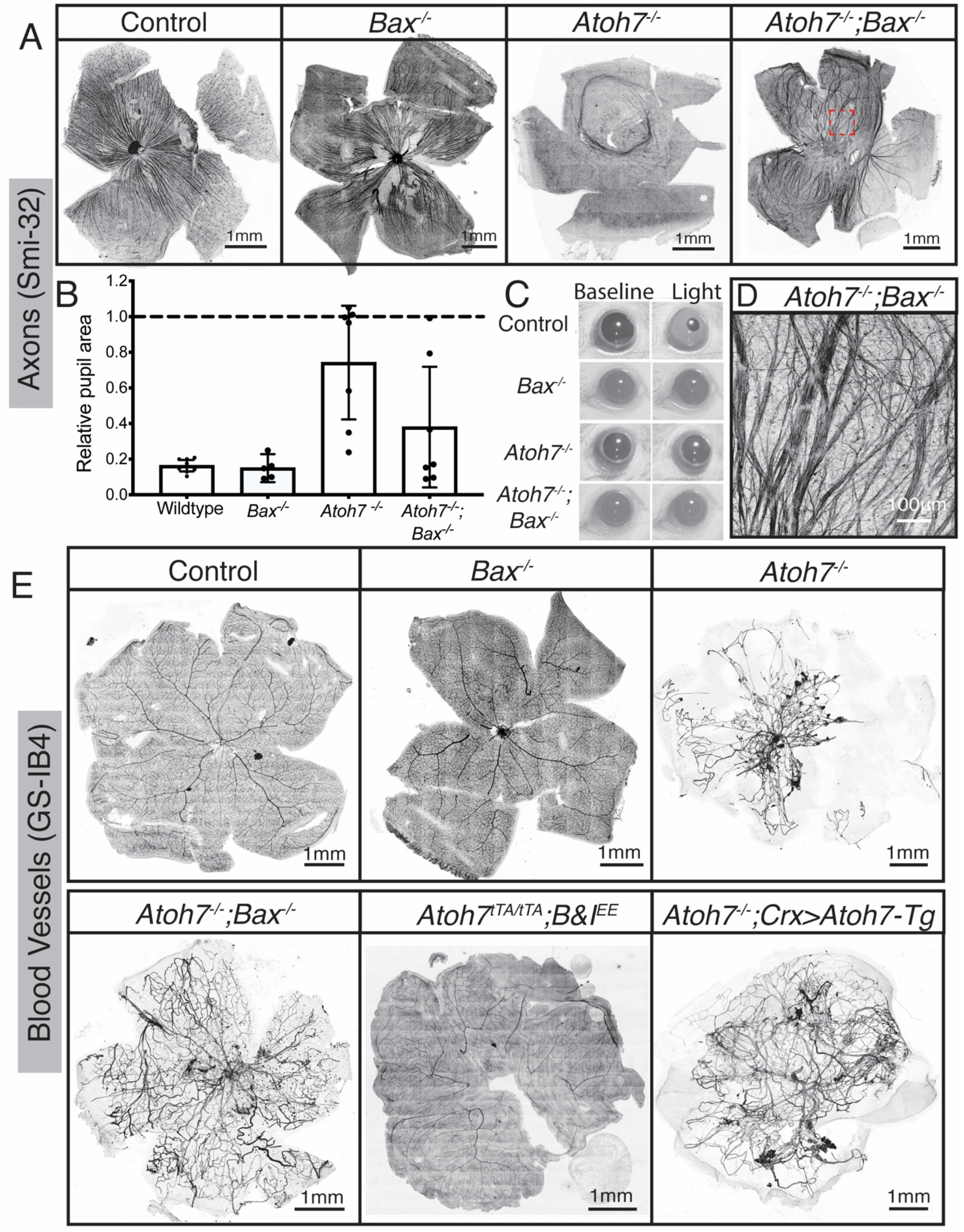
RGC axon guidance and retinal vasculature development requires *Atoh7*-dependent RGCs. (A,D) Smi-32 labels a subset of RGCs and their axons in an adult wildtype retina. In *Atoh7*^*−/−*^ mice, the Smi-32 positive RGCs have axon guidance deficits. In *Atoh7*^*−/−*^*;Bax*^*−/−*^ mice, RGCs have severe axon guidance deficits. Highlighted region (A - *Atoh7*^*−/−*^*;Bax*^*−/−*^) is magnified in (D). (B,C) Using the contralateral pupillary light response as a readout of retina to brain connection allows the appreciation that the severe axon guidance deficits allow for some connection to the brain of the RGCs in the *Atoh7*^*−/−*^ or *Atoh7*^*−/−*^*;Bax*^*−/−*^ retinas. (E) It has been previously reported that the hyaloid vasculature fails to regress in *Atoh7*^*−/−*^ mice, thought to be due to lack of RGCs, however when the RGC numbers are rescued, in *Atoh7*^*−/−*^*;Bax*^*−/−*^mice, the hyaloid vasculature fails to regress. However, *Atoh7* is not necessary for the hyaloid regression and retinal vasculature development, seen using *Atoh7*^*tTA/tTA*^*;B&I*^*EE*^ mice, which was previously seen to rescue all of the *Atoh7* null phenotypes. When *Atoh7* is rescued using the *Crx>Atoh7* transgene on the *Atoh7* null background, the optic nerve and 12% of RGCS are rescued (Prasov and Glaser, 2012), but the hyaloid vasculature does not regress.

Previous studies have observed a lack or massive reduction in physical or functional connection to the brain in *Atoh7*^*−/−*^ mice (Brown et al., 2001; Brzezinski et al., 2005; Triplett et al., 2011; Wee et al., 2002). Consistent with the failure of mutant RGCs to correctly target the optic nerve, we observe severe disruptions in behavioral responses to light in *Atoh7*^*−/−*^*;Bax*^*−/−*^ mice that are essentially indistinguishable from those seen in *Atoh7*^*−/−*^ mice. *Atoh7*^*−/−*^*;Bax*^*−/−*^ mice show no detectable optokinetic response (Supplemental Figure 7A), and show no visual cue-dependent reduction in escape time during successive trials of the Morris water maze (Supplemental Figure 7B). *Opn4:Tau-lacZ* knock-in mice, which visualize the axonal projections of M1 ipRGCs, show no detectable signal in the brain (Supplemental Figure 7C) (Hattar et al., 2002). Intraocular injection of fluorescently-labeled cholera toxin beta, which visualizes RGC axonal terminals (Bedont et al., 2014), likewise shows no brain labeling in both *Atoh7*^*−/−*^*;Bax*^*−/−*^ and *Atoh7^−/−^ mice* (Supplemental Figure 7D). However, while the contralateral PLR is significantly reduced compared to wildtype in both the Atoh7^*−/−*^ and *Atoh7*^*−/−*^*;Bax*^*−/−*^ mice, it is nonetheless detectable, indicating that a small number of RGC axons target the olivary pretectal nucleus in *Atoh7*^*−/−*^*;Bax*^*−/−*^ mice, although we are unable to detect these using standard techniques (Figure 3B,C, Supplemental Figure 5 A-C).

### Retinal vasculature development is disrupted in the absence of *Atoh7*

In both mice and humans, loss of *Atoh7* expression results in persistence of the hyaloid vasculature (Edwards et al., 2012; Ghiasvand et al., 2011; Kondo et al., 2016; Prasov et al., 2012). The persistence of the hyaloid vasculature in *Atoh7*^*−/−*^ retinas until P14 was previously observed. We likewise observe persistence of the hyaloid vasculature into adulthood in *Atoh7*^*−/−*^ retinas (Figure 3E), which is observed even in one year old mice (data not shown). Surprisingly, even with the rescue of a majority of Brn3a RGCs in *Atoh7*^*−/−*^*;Bax*^*−/−*^ animals, the hyaloid vasculature still fails to regress (Figure 3E). Likewise, *Crx>Atoh7;Atoh7*^*−/−*^ mice, in which *Atoh7* is misexpressed in photoreceptor precursors, also fail to induce hyaloid regression (Figure 3E). This is in sharp contrast to the rescued vascular phenotype observed in *Atoh7*^*tTA/tTA*^*;B&I-EE* mice, when *Brn3b* & *Isl1* are ectopically expressed from the endogenous *Atoh7* locus in a *Atoh7*-deficient mouse using the tet-off system (Wu et al., 2015)(Figure 3E), and implies that *Brn3b* and *Isl1* may activate expression of secreted factors that drive vascular regression.

### RGC differentiation is delayed in the absence of *Atoh7*

In order to examine potential differences in RGC development within *Atoh7*^*−/−*^ and *Atoh7*^*−/−*^*;Bax*^*−/−*^ compared to wildtype and *Bax*^*−/−*^ control animals, we next performed single-cell RNA-sequencing on *Bax*^*−/−*^, *Atoh7*^*−/−*^, and *Atoh7*^*−/−*^*;Bax*^*−/−*^ retinas to more comprehensively profile changes in cell type specification and global transcriptional differences across the genotypes. We profiled 67,050 E14.5 retinal cells from *Bax*^*−/−*^, *Atoh7*^*−/−*,^ and *Atoh7*^*−/−*^*;Bax*^*−/−*^ mice, and aggregated the datasets with 26,078 age-matched wildtype retinal cells from previous studies (Clark et al., 2019) (Figure 4A,B, Supplemental Figure 8). Consistent with the previous single-cell studies of E14 mouse retinal development (Clark et al., 2019; Giudice et al., 2019), we observed a continuous manifold of cells, from primary RPCs to neurogenic RPCs, leading to three separate differentiation trajectories that give rise to RGCs, cones, and amacrine/horizontal cells, respectively (Figure 4A-B; Supplemental Figure 9A). Analysis of the cell type proportions across the genotypes revealed a depletion of RGCs in both *Atoh7* mutant samples (*Atoh7*^*−/−*^ and *Atoh7*^*−/−*^*;Bax*^*−/−*^) compared to both wildtype and *Bax*^*−/−*^ controls (Figure 4C-D, Supplemental Figure 8E-F). This reduction in the number of RGCs included a compensatory increase in cone photoreceptors and neurogenic RPCs, consistent previous findings (Figure 4C-D, Supplemental Figure 8E-F) (Brown et al., 2001; Brzezinski et al., 2005; Hufnagel et al., 2010; Wang et al., 2001).

**Figure 4.**
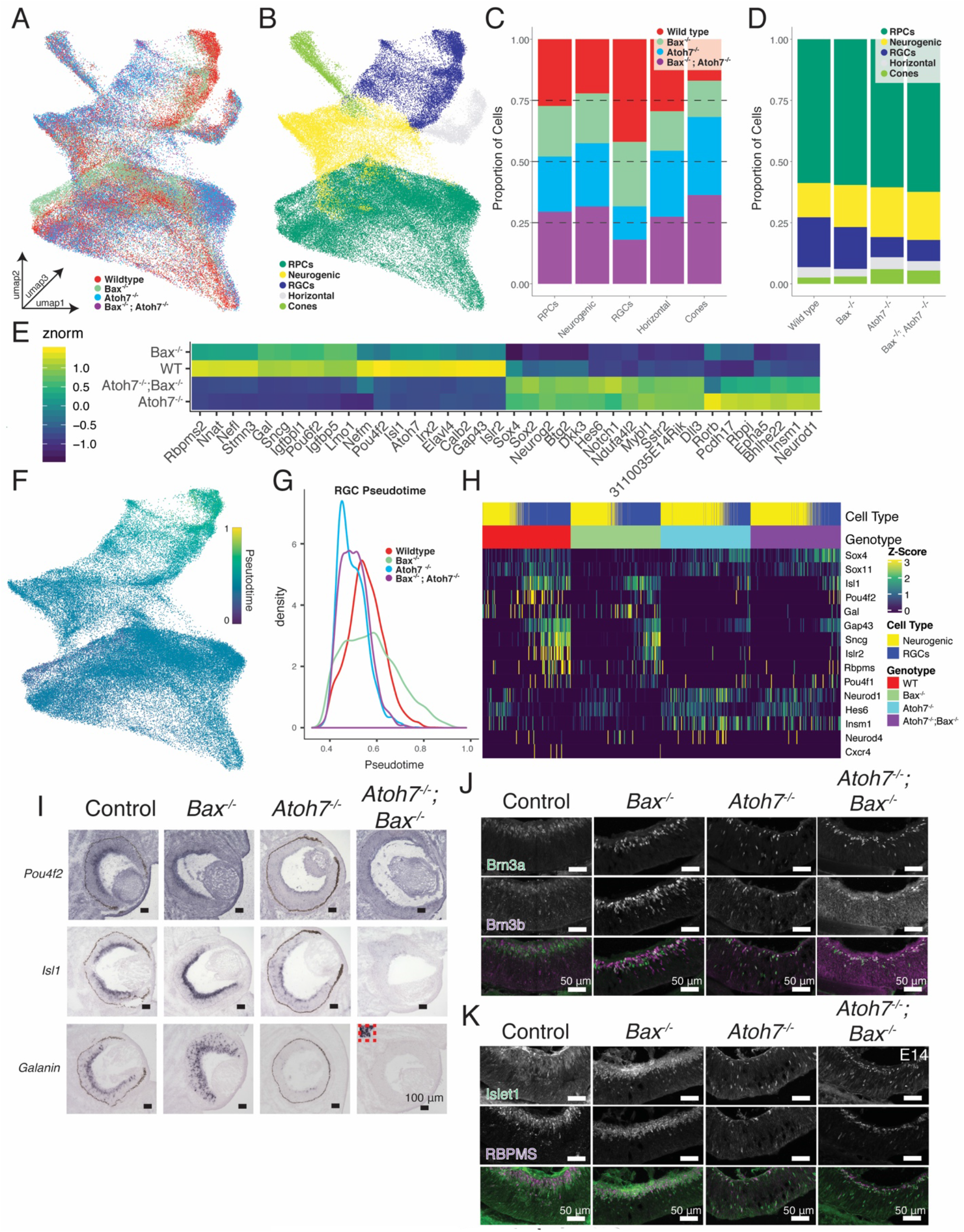
Single-cell analysis of E14.5 mutant retinas and gene detection in E14 retinas. (A,B) UMAP dimension reduction of aggregated E14.5 single cell dataset colored by (A) genotype and (B) annotated cell-type. (C) Proportions of cell types derived from each genotype. (D) Proportions of annotated cell types within each genotype. (E) Heatmap of differentially expressed transcripts across control and Atoh7 knockout (*Atoh7*^*−/−*^ or *Atoh7*^*−/−*^*;Bax*^*−/−*^) neurogenic and RGCs. (F) UMAP dimension reduction of cells colored by Scanpy pseudotime values. (G) Density of retinal ganglion cells along pseudotime by genotype. (H) Heatmap displaying differentially expressed transcripts across the interaction of pseudotime and genotype. Cells are ordered by pseudotime within each genotype. (I) Chromogenic *in situ* hybridization detecting RNA transcripts of genes from (H). Insert, depicted by a red dotted line, *Atoh7*^*−/−*^*;Bax*^*−/−*^ mice in (H) show robust galanin signal in a region outside the retina, but minimal signal in the retina (J-K). Immunohistochemistry detecting (J) RGC-specific markers: BRN3A and BRN3B and (K) pan-RGC markers: ISL1 and RBPMS in E14 retina from each genotype. Scale bar, 50 μm. Abbreviations: RPCs, retinal progenitor cells; RGCs, retinal ganglion cells.

**Figure 5.**
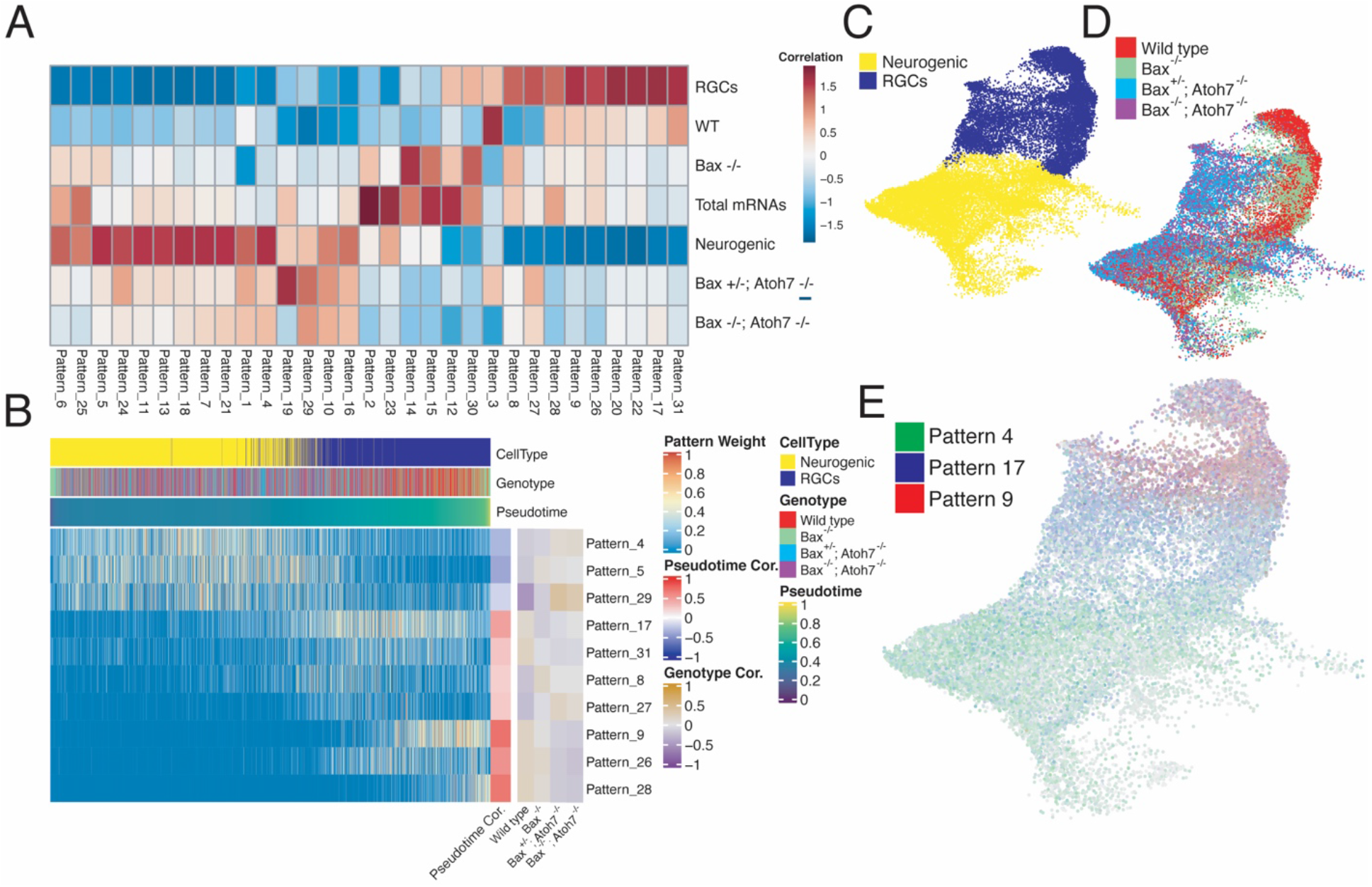
scCoGAPs analysis of single-cell dataset and RGC population changes in E12.5 retinas show a developmental delay in *Atoh7*^*−/−*^*;Bax*^*−/−*^. mutants. (A) Heatmap showing the correlation between scCoGAPS pattern and cellular features. (B) Heatmap of pattern weights within individual cells ordered by pseudotime. Pattern correlations with both pseudotime and each genotype are displayed at the right. (C-E) UMAP embedding of single-cell dataset used for scCoGAPS and colored by (C) celltype or (D) genotype. (E) UMAP embedding of dataset and colored by pattern weights of scCOGAPS patterns 4, 17 and 9, displaying progressive pattern usage across RGC development.

Using the scRNA-seq data, we next assessed the differential gene expression within neurogenic RPCs and RGCs across control -- WT and *Bax*^*−/−*^ -- and *Atoh7*- deficient -- *Atoh7*^*−/−*^ and *Atoh7*^*−/−*^*;Bax*^*−/−*^ -- samples. Using strict differential expression cutoffs (q-value < 1e-300), we identified 230 *Atoh7*-dependent differentially expressed transcripts within neurogenic RPCs and RGCs (Supplemental Table 1). Genes enriched within control samples (Figure 4E) highlighted many known factors in the specification and differentiation of RGCs, including *Pou4f2* (*Brn3b*), *Isl1*, *Pou6f2* (*Rpf-1*), *Elavl4*, *Gap43*, and *Irx2* (Choy et al., 2010; Ekström and Johansson, 2003; Kruger et al., 1998; Zhou et al., 1996) (Supplemental Figure 9B). Conversely, differentially expressed transcripts enriched in the *Atoh7* knockout samples (Figure 4E) were enriched for genes involved in the Notch-signaling pathway -- *Rbpj, Dll3, Notch1,* and *Hes6* -- and for transcripts enriched in neurogenic cells and photoreceptor precursors during retinal development -- *Btg2, Neurog2, Bhlhe22, Insm1, Neurod1, Mybl1, Sstr2, 3110035E14Rik* (Clark et al., 2019; Supplemental Figure 9B).

*Atoh7-*deficient RGCs also show dramatically reduced expression of genes known to regulate axon guidance including *Islr2*, which has been found to control RGC axon fasciculation, as well as axon guidance at the optic chiasm (Panza et al., 2015). Of particular interest are the observed increases in *Rbpj* and *3110035E14Rik* expressions in *Atoh7*^*−/−*^ and *Atoh7*^*−/−*^*;Bax*^*−/−*^ samples. *Rpbj* is an upstream regulator of *Atoh7* expression (Miesfeld et al., 2018b), and *3110035E14Rik* (*Vxn*), which functions similarly to Atoh7 by promoting retinal neurogenesis and early retinal cell fate specification (Moore et al; 2018). The increased expression of transcripts that show enriched expression in neurogenic cells and photoreceptor precursors is consistent with the increase in generation of cone photoreceptor precursors seen in an *Atoh7-*deficient retina. Combined with a recovery in the specification of RGC numbers in adult *Atoh7*^*−/−*^*;Bax*^*−/−*^ animals, these data suggest that loss of *Atoh7* expression leads to an increase in expression of genes specific to neurogenic RPCs at the expense of RGC-enriched transcripts, results consistent with a developmental delay.

In order to further assess the degree of an RGC-specific developmental delay we performed pseudotemporal analyses using Scanpy (Wolf et al., 2018)(Figure 4F). We observed a bias of the *Atoh7*-deficient cells within early pseudotime stages during RGC differentiation. Both wildtype and *Bax*^*−/−*^ control cells had a broader distribution of cells across pseudotime, results consistent with a failure of maturation or developmental delay of RGC specification in *Atoh7*-deficient RGCs (Figure 4G). Differential expression analysis assessing the changes in gene expression across the interaction of pseudotime and genotypes revealed significant genotypic differences across RGC development (Figure 4H; Supplemental Table 2).

In both *Atoh7*^*−/−*^ and *Atoh7*^*−/−*^*;Bax*^*−/−*^ samples, we observed a reduction of expression in many genes enriched within mature RGCs -- *Pou4f2, Gap43, Sncg, and Isl1*. We likewise observed reduced expression of a subset of genes in neurogenic RPCs, including *Gal*. Increased expression of other genes predominantly expressed in neurogenic RPCs – including *Neurod1, Insm1, Neurod4, Hes6, Onecut1, Onecut2,* and *Sox4 --* is observed in both *Atoh7*-deficient neurogenic RPCs and RGCs compared to controls (Figure 4H). This implies that loss of function of *Atoh7* may delay differentiation of RGCs from neurogenic RPCs. These results overall, closely match those obtained from scRNA-Seq-based analysis of *Atoh7*^*−/−*^ retina conducted at E13.5 (Wu et al., 2020).

We next performed *in situ* hybridization to examine changes in global transcript expression within the developing retina. RNA transcript expression was detected at E14.5, at which point most RGCs are specified (Sidman, 1960; Young, 1985), and we observed decreased expression of *Pou4f2 (Brn3b), Isl1* and *Gal*, in both *Atoh7*^*−/−*^ and *Atoh7*^*−/−*^*;Bax*^*−/−*^ mice, as determined by chromogenic *in situ* hybridization (Figure 4I). Immunostaining of E14 retinas confirm a reduction in the number of cells immunopositive for Brn3a (Pou4f1) and Brn3b (Pou4f2) (Figure 4J), as well as the pan-RGC markers RBPMS and Isl1 (Figure 4K), in the developing ganglion cell layer of *Atoh7-*deficient retinas. At E12.5 we observed a marked decrease in both overall RGC density and RGC number (Supplemental Figure 3). Together these results suggest that loss of function of *Atoh7* delays RGC differentiation, and leads to an accumulation of neurogenic RPCs.

To further identify patterns of temporal changes in gene expression across RGC genesis between *Atoh7* mutant and control retinas, we performed the non-negative matrix factorization technique scCoGAPS (Stein-O’Brien et al., 2019). Implementation of scCoGAPS parses the gene expression into groups (‘patterns’) based on gene expression profiles and without *a priori*, literature-based knowledge of gene interactions. Using 5235 highly variable genes across the 29,182 neurogenic RPCs and differentiating RGCs, we identified 31 patterns of gene expression (Supplemental Figure 10). These patterns correlated with both neurogenic RPCs – Patterns 6, 25, 5, 24, 11, 13, 18, 7, 21, 1, and 4 – and RGC – Patterns 8, 27, 28, 9, 26, 20, 22, 17, 31 – cell type annotations and highlighted temporal changes in gene expression as assessed through pseudotime analyses. Individual patterns - patterns 4, 5, and 29 - were highly correlated with neurogenic cell annotations and *Atoh7*^*−/−*^ and *Atoh7*^*−/−*^*;Bax*^*−/−*^ genotypes, consistent with a temporal delay in RGC specification, increased number of neurogenic cells, or failure of cell type specification of neurogenic RPCs as a consequence of lost *Atoh7* expression. Genes driving pattern 5 include genes enriched in early-stage neurogenic RPCs, *Gadd45a* and *Sox11* (Clark et al., 2019).

Conversely, patterns highly correlated with RGC cell type annotations and pseudotime – Patterns 9, 26, and 28 – had high correlation with control samples. The most highly weighted genes in Patterns 9 and 26 are *Gap43* and *Igfbpl1*, respectively, which have been implicated in RGC axonal growth (Guo et al., 2018; Zhang et al., 2000). Pattern 28 highlights cells towards the end of the RGC trajectory and is largely driven by Sncg; a transcript enriched in most RGCs in the adult mouse retina (Soto et al., 2008). The association of neurogenic patterns with *Atoh7*^*−/−*^ mutant retinas versus those that highlight RGC differentiation and maturation patterns with control retinas further support a developmental delay in mutant RGCs. Analysis of pattern marker expression across the genotypes (Supplemental Figure 10) highlights both the temporal delay and global changes in gene expression across the *Atoh7*^*−/−*^ mutant retinas compared to controls.

Recent studies have comprehensively profiled RGC subtype diversity in the mouse retina (Rheaume et al., 2018; Tran et al., 2019). However, these studies did not characterize either the birthdates of individual RGC subtypes, or the transcriptional networks controlling RGC subtype specification. Our combined data supports evidence of both *Atoh7-*dependent and independent populations of RGCs. The delay in RGC maturation and the failure of optic nerve formation seen in *Atoh7-*deficient retinas suggest that the earliest pathfinding RGCs are *Atoh7*-dependent. We examined expression of markers of mature RGC subtypes (Tran et al., 2019) within the developing retina and correlated expression of the transcripts with RGC pseudotime (Supplemental Figure 11) as many of the mature RGC subtype markers are not specific to RGCs. We detected expression of selective markers for a fraction of mature RGC subtypes within the E14.5 scRNA-seq dataset. Of transcripts in which readily detectable expression was observed, many – including *Igfbp4, Foxp1, Stxbp6, Bhlhe22, Penk* – also display enriched expression in primary or neurogenic RPCs. Expression of some markers of RGC development and maturation – *Ebf3, Pou4f1, Pou4f2, Prdm8* and *Slc17a6* – correlated well with pseudotemporal ordering and were depleted in *Atoh7*-deficient RGCs. However, a limited number of RGC subtype markers, including *Irx3, Calb2,* and *Tac1* were largely absent from *Atoh7* mutant RGCs.

Our data suggest the existence of *Atoh7*-dependent factors that promote both RGC survival and pathfinding, and also drive hyaloid vascular regression. These are likely derived from *Atoh7*-expressing neurogenic RPCs and/or RGCs. Factors mediating hyaloid regression are likely to be secreted, while those regulating RGC survival could also potentially act through contact-mediated signaling. Few annotated secreted proteins show clear *Atoh7*-dependent expression in our scRNA-Seq dataset. The neuropeptide galanin *(Gal*), which is strongly expressed in both neurogenic RPCs and RGCs, was by far the most differentially expressed secreted factor in *Atoh7*-deficient mice (Figure 4E,I, Supplemental Table 1). Galanin has been implicated in promoting the survival of neural precursors (Cordero-Llana et al., 2014; Holmes et al., 2000), however, *Gal*-deficient animals showed no differences in either the hyaloid vasculature regression or RGC density, as compared to the control animals (Supplemental Figure 12).

## Discussion

It is broadly accepted that *Atoh7* acts in RPCs as a competence factor that is essential for RGC specification (Brzezinski et al., 2012; Mu et al., 2005; Yang et al., 2003; Baker and Brown, 2018). In this study, though, we show that specification of the great majority of RGCs occurs normally even in the absence of *Atoh7*. While RGC specification can occur independent of *Atoh7*, *Atoh7* function is required to maintain RGC survival and proper targeting of RGC axons to the optic nerve head. Following disruption of both *Atoh7* and *Bax*, we observe only a 20% reduction in the number of RGCs relative to *Bax*-deficient controls. This compares to a greater than 95% reduction in RGC numbers in *Atoh7* mutants relative to wildtype controls. Although RGCs in *Atoh7*^*−/−*^*;Bax*^*−/−*^ retinas show severe defects in targeting the optic nerve head, they respond robustly to photoreceptor stimulation. The presence of specified RGCs in the absence of Atoh7 helps explain long-standing, puzzling observations: 1) 45% of RGCs are not derived from *Atoh7*-expressing progenitors; 2) molecular markers of RGCs are observed at considerably higher levels during early stages of retinal development than in adults in Atoh7-deficient retinas; and 3) the marked increase in apoptosis in the ganglion cell layer that occurs in the absence of Atoh7 (Brzezinski et al., 2012; Feng et al., 2010; Prasov and Glaser, 2012). Previous studies have implicated Atoh7 as a direct upstream regulator of the essential RGC transcription factors *Brn3a*, *Brn3b*, and *Isl1*. Supporting this, an *Atoh7* hierarchy of RGC determinants, studies in which Brn3b and Islet1 were inserted in place of the Atoh7 coding sequence observed a complete rescue of normal RGC development (Wu et al., 2015). Our studies, however, indicate that when apoptosis is inhibited, Brn3a, Brn3b, Islet1, and Rbpms expression is induced at near normal levels within RGCs in both E14 and adult retinas, regardless of Atoh7 expression. Therefore, we posit that factor(s) in addition to Atoh7 activate expression of these genes.

The molecular mechanisms by which *Atoh7* controls axonal guidance and cell survival remain unclear. One hypothesis is that RGC axonal guidance and cell survival are linked and that loss of Atoh7 expression fails to initiate expression of target-derived trophic cues. Because other studies have failed to detect substantial numbers of RGCs in *Atoh7*^*−/−*^ mice at P0 (Brown et al., 2001; Feng et al., 2010; Prasov and Glaser, 2012; Wang et al., 2001) and target derived neurotrophic factors regulate apoptosis of RGCs between P0 and P12, we feel that Atoh7-induced cell survival is functioning at earlier time points than target-derived trophic signals. Additionally, as we did not observe RGC rescue within *Atoh7*^*Cre/Cre*^*;Bax*^*fl/fl*^ mice, we suggest that immature RGCs rapidly degenerate as the result of the lack of an *Atoh7*-dependent survival factor. This wave of developmental apoptosis has been observed previously, but the underlying molecular mechanisms are unknown (Farah and Easter, 2005; Frade et al., 1997; Ogilvie et al., 1998; Péquignot et al., 2003; Rodríguez-Gallardo et al., 2005; Strom and Williams, 1998; Valenciano et al., 2009). We hypothesize that this unknown factor or factors must be produced by either *Atoh7*-expressing neurogenic RPCs or RGCs derived from these cells.

Our data also reveal a marked delay in formation of RGCs from neurogenic RPCs in the absence of *Atoh7*. This is consistent with results from others that in *Atoh7*^*−/−*^ retina, RGC formation is delayed by at least a day (Le et al., 2006; Prasov and Glaser, 2012). When *Atoh7*-dependent RGCs are rescued later in development, as seen in targeted mutants in which *Atoh7* is expressed from the endogenous *Crx* locus, the hyaloid vasculature regression was not rescued, even though a modest rescue of RGC formation is observed (Prasov and Glaser, 2012). In zebrafish retina, consistent with this result, loss of function of *atoh7* in early-stage RPCs disrupts correct targeting of axons in later-born RGCs to the optic nerve (Pittman et al., 2008).

Since *Atoh7* was previously thought to be a master transcriptional regulator of RGC specification, strategies aimed at targeted differentiation of RGCs for therapeutic purposes have focused on using forced expression of *Atoh7*. We, however, now appreciate RGC specification to be a far more complicated process. Although ectopic expression of *Atoh7* activates expression of RGC-specific genes in cultured retinal progenitor cells (Yao et al., 2007), induced pluripotent stem cells (Chen et al., 2010), and Müller glia derived retinal stem cells (Song et al., 2015, 2014), it is nonetheless typically not sufficient to drive these cells to become RGCs. This study sheds light on why this may be the case.

These findings demonstrate that additional factors act in parallel to *Atoh7* to control RGC specification. While multiple other transcription factors have been reported to regulate RGC specification -- including *Neurod1*, *Sox4*, and *Onecut2* (Jiang et al., 2013; Mao et al., 2013; Sapkota et al., 2014) -- these factors are unable to individually activate expression of *Brn3b* and *Isl1*. In *Atoh7*^*−/−*^*;Bax*^*−/−*^ RGCs, however, we observe substantially increased expression of each of these transcription factors (Figure 4E; Supplemental Figure 10), suggesting the possibility that these factors, among others, may compensate for the loss of *Atoh7*. The identity of the non-cell-autonomous cues by which *Atoh7* regulates RGC survival, axon guidance, and hyaloid vasculature regression remain unknown. Further identification of the mechanisms regulating the interplay of intrinsic and extrinsic signals on RGC specification, survival, and maturation will help guide the future design of therapies aimed at maintaining and/or replacing damaged/lost RGCs.

## Acknowledgements

We thank Xiuqian Mu and Fuguo Wu for providing *Atoh7*^*tTA/tTA*^*;B&I-EE* mice, Lin Gan for providing *Atoh7*^*Cre*^ mice, Nadean Brown for technical advice, Haiping Hao and Hopkins Transcriptomics and Deep-Sequencing Core for assistance with scRNA-Seq, and Wendy Yap for comments on the manuscript. This work was supported by NIH Grants R01EY020560 (SB), R00EY27844 (BSC), EY19497 (TG), GM076430 and EY027202 (SH & HZ), EY021372 (JHS), and F32 EY022543 (TS), the intramural research fund at the National Institute of Mental Health (MH002964) (SH), and an unrestricted grant to the Department of Ophthalmology and Visual Sciences from Research to Prevent Blindness (BSC).

## Supplemental Methods

### Morris water maze

Vision was tested using a cued Morris water maze (Morris, 1984). A 150cm diameter pool was filled with water made opaque by mixing in non-toxic white tempera paint. A platform (“island”, not visible from the surface of the water) was placed in one quadrant with a black and white striped 50 mL conical tube on top as a “flag”. Each of six animals in a group (either *Atoh7*^*-/+*^, *Atoh7*^*−/−*^, or *Atoh7^−/−^;Bax^−/−^)* was allowed a 90 second trial in which to swim and visually locate the island. If the animal found the island, it was allowed to remain on the island for 60 seconds before being removed from the pool. If the animal did not locate the island, it was placed at the edge of the island, allowed to climb on, and remain on the island for 60 seconds to reinforce the island as a safe spot. Following the first trial, the island was moved to a different quadrant and animals were tested again. In total, four trials were completed with the island in four different locations.

### Visual tracking

Visual acuity was assessed using OptoMotry (Cerebral Mechanics). Animals were placed on a platform in the viewing chamber and allowed to adapt for 10 minutes prior to testing. Testing was done for five minutes, during which the position of the animal’s head was monitored to assess visual tracking. Following each tracking event, the direction and width of the grating was changed, eventually narrowing down to smaller ranges of acuity.

### X-Gal Staining

To examine ipRGCs and projections in animals with one allele of Opn4^tauLacZ/+^, adult animals were transcardially perfused with 10 mL PBS followed by 15 mL of 4% PFA. Brains were dissected and cryoprotected in 30% sucrose in PBS until the tissue had sunk in the solution (kept at 4°C) and then frozen in OCT. Cryosections (50 μm) were mounted on slides and allowed to dry up to 8 hours at room temperature. All X-Gal staining solutions were prepared according to (Mombaerts et al., 1996). Slides were rehydrated in Buffer B for 10 minutes, then immersed in Buffer C (with potassium ferricyanide, potassium ferrocyanide and 100 mg/mL X-Gal in N-N’ dimethyl formamide (DMF)) for 2 days in a dark container. On the third day, fresh Buffer C was added to the slides which were allowed to sit overnight. Following staining, slides were washed 3 times for 5 minutes in PBS, counterstained in Nuclear FastRed (diluted 1:5 in water) for 10 minutes, washed in 70% ethanol, and mounted in glycerol. Retinas were dissected in phosphate buffer and placed in X-Gal Buffer B for 10 minutes, then Buffer C for 2 days at room temperature in a dark container. On the third day, fresh Buffer C was added, and retinas were stained for an additional day. Following staining, retinas were washed 3 times for 5 minutes in PBS, washed in 70% ethanol, and flat mounted in glycerol.

### CtB sectioning and visualization

Pulled glass needles were filled via capillary action with roughly 5 μL of Alexa 488 conjugated Cholera toxin b subunit (CTb; Invitrogen) diluted to 5mg/mL in dH20. Filled needles were affixed to a Harvard Apparatus FIL-190 picospritzer. For injection, adult animals were anaesthetized with 25ul/g body weight Avertin (1.25% tri-bromo-ethanol and 2.5% tertiary-amyl alcohol in dH20) and tested for alertness via toe pinch. Gentle pressure was applied behind the eye to raise it, the needles were aimed just above the skin margin, and CTb was injected in the intraocular space. Animals were allowed to recover and maintained for three days post-injection to allow the tracer to reach the distal axons tips. Following the three-day survival period, animals were transcardially perfused with 15 mL of PBS, followed by 25 mL of cold 4% PFA. Brains were dissected after perfusion and post-fixed in 4% PFA for 1 hour, then cryoprotected in 30% sucrose. Brains were frozen in OCT and sectioned at 50 μm in a cryostat. Sections were dried overnight, fixed in 4% PFA for 10 min, mounted in VectaShield (Vector Labs)+1.5 mg/mL DAPI and imaged on a Zeiss Imager Ml upright epifluorescence microscope (AxioVision software).

## Supplemental Figure Legends

**Supplemental Table 1.** 230 differentially expressed transcripts (q-value < 1e-300) between control (control and *Bax*^*−/−*^) versus *Atoh7*-deficient *(Atoh7*^*−/−*^, or *Atoh7*^*−/−*^*;Bax*^*−/−*^) neurogenic and retinal ganglion cells.

**Supplemental Table 2.** Differentially expressed transcripts (q-value < 1e-5) across the interaction of pseudotime and genotype.

**Supplemental Figure 1.**
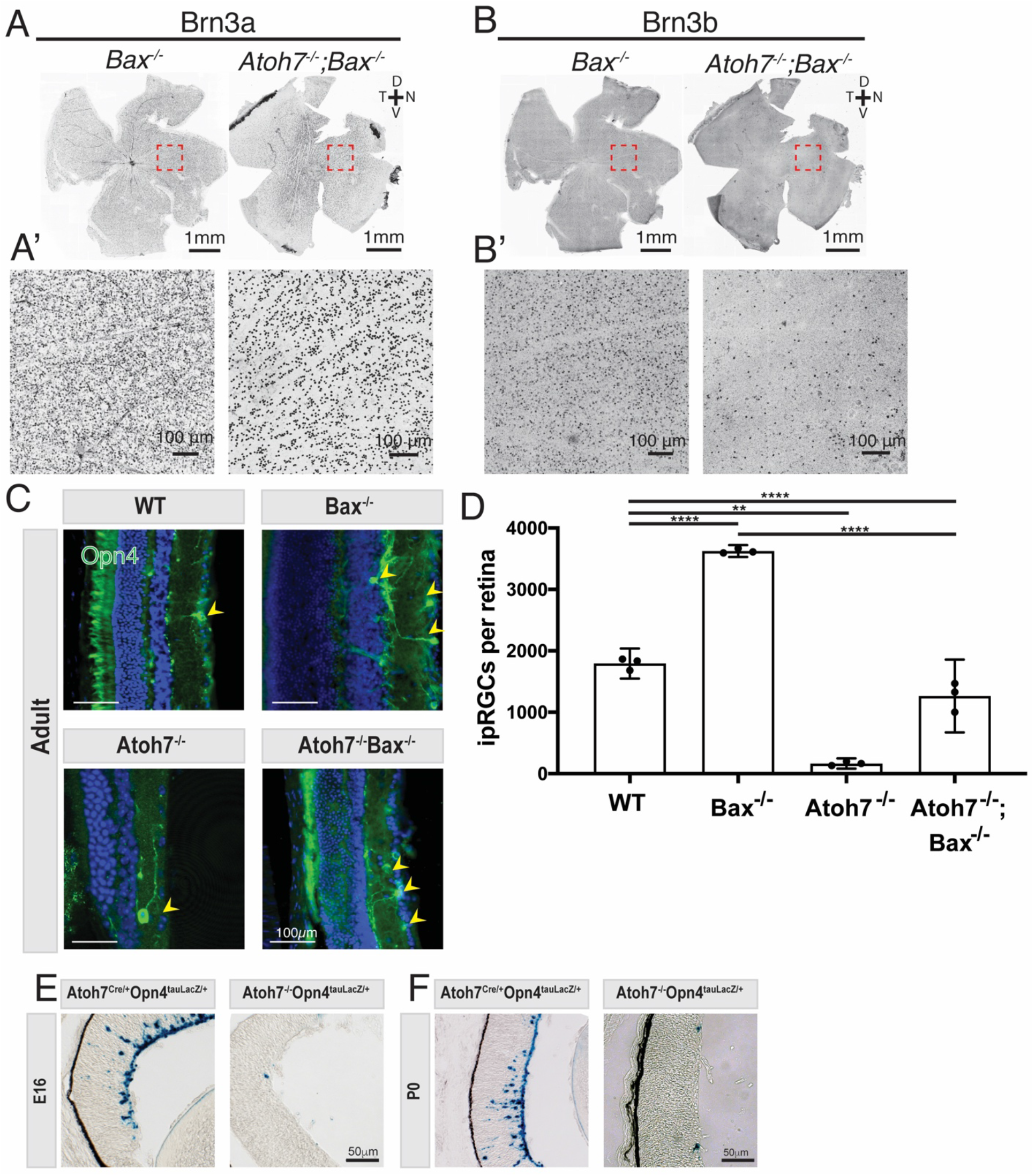
(A,B) Whole retinal dissection showing the presence of Brn3a and Brn3b positive RGCs respectively. Notice the orientation of the retinas as marked by the orientation rose. Highlighted regions (A,B) are magnified in (A’,B’). (C,D,E,F) IpRGCs are partially rescued by preventing apoptosis. (C,D) To determine if ipRGCs were undergoing apoptosis in the Atoh7^−/−^, ipRGC numbers were examined in Atoh7^−/−^ and *Atoh7*^*−/−*^*;Bax*^*−/−*^. ipRGCs in the *Atoh7*^*−/−*^*;Bax*^*−/−*^ were rescued to 60% of wild type levels. *Bax*^*−/−*^ showed a 202±9% increase in ipRGCs compared to controls, similar to what has been reported for other RGCs in the *Bax*^*−/−*^ knockout. (E,F) To determine if ipRGCs were generated in development in Atoh7^−/−^ mice, I examined E16 and P0 timepoints using the Opn4^tauLacZ^ marker. At both E16 and P0, a 90% deficit of ipRGCs was observed. Mean ± 95% confidence intervals. Statistical significance tested by one-way ANOVA with Tukey’s post test for multiple comparisons ** p=0.0052, **** p < 0.0001.

**Supplemental Figure 2.**
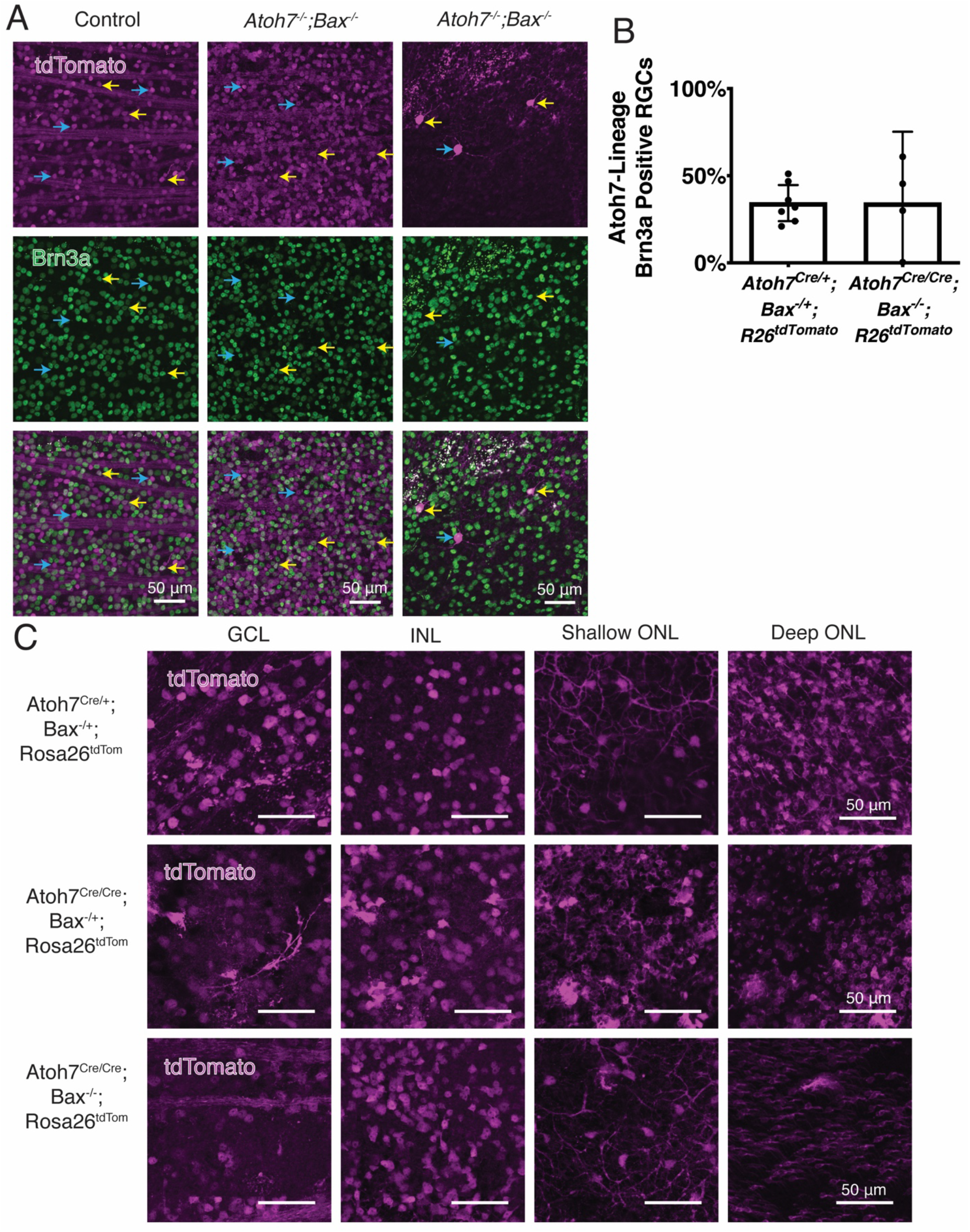
Atoh7^Cre^;R26^tdTomato^ lineage labeling in detail. (A,B) A similar percent of RGCs are of the Atoh7 lineage in both wildtype and *Atoh7*^*−/−*^*;Bax*^*−/−*^ mice. Yellow arrows highlight Brn3a+ and Atoh7-expressing RPC-derived RGCs while blue arrows highlight non-Brn3a+ cells in the GCL (ganglion cell layer). (C) Atoh7^Cre^;R26^tdTomato^ labels cells of most cell types in the retina in all genotypes examined. Abbreviations: GCL, INL (inner nuclear layer), ONL (outer nuclear layer).

**Supplemental Figure 3.**
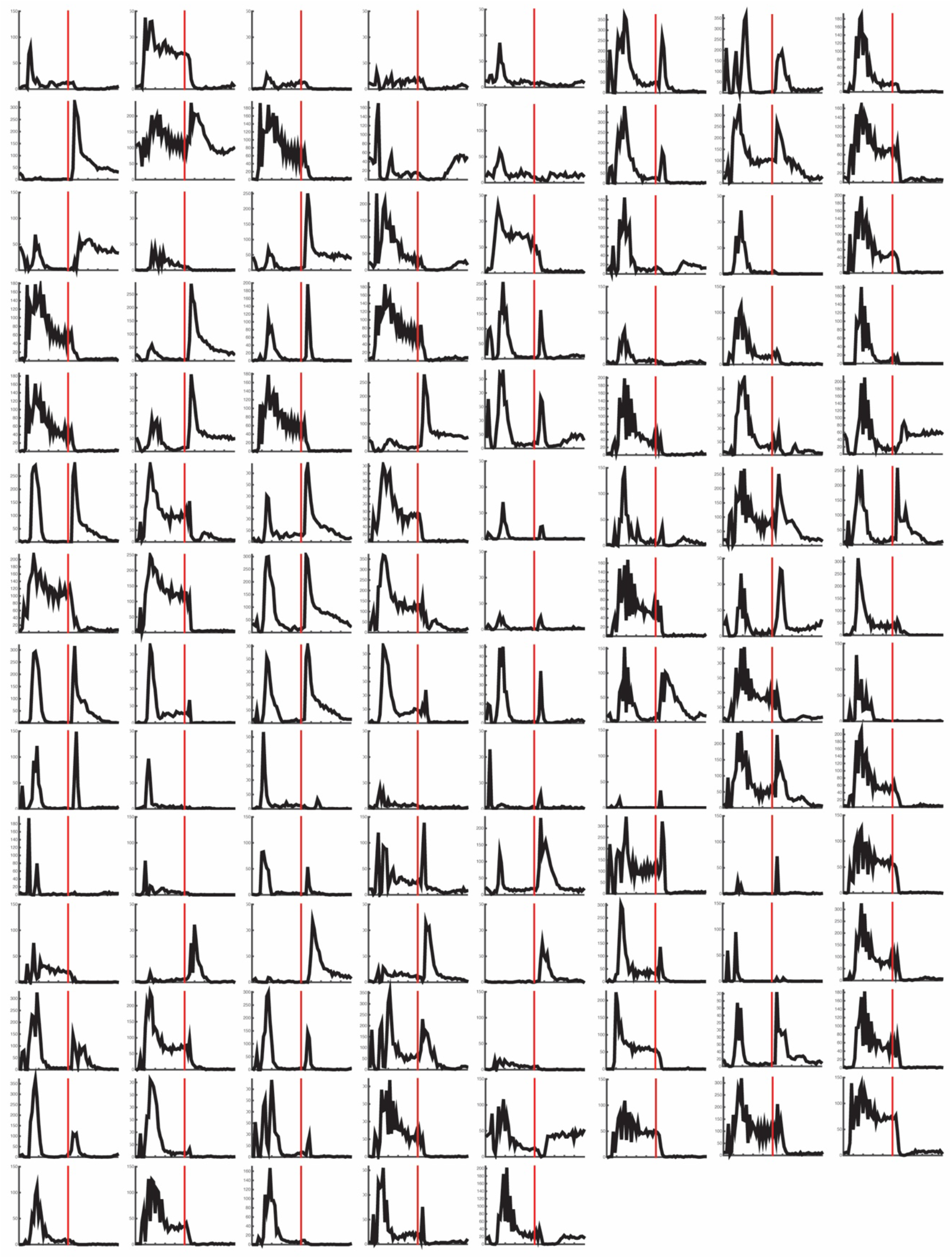
During early development, E12.5, *Atoh7*^*−/−*^*;Bax*^*−/−*^ mice have both fewer Brn3a positive RGCs and mature RGCs. (A) Pax2 antibody staining was used to help find the central retina to make sure the earliest born RGCs in the central retina were fully sampled. (B) Quantification of the retinal images with the black bars representing density and the red bar plotted on the right side representing the percent of mature RGCs. Mean ± 95% confidence intervals. Statistical Significance tested by unpaired t test ** p = 0.0099 **** p < 0.0001. Abbreviations: RGCs, retinal ganglion cells; GCL, ganglion cell layer.

**Supplemental Figure 4.**
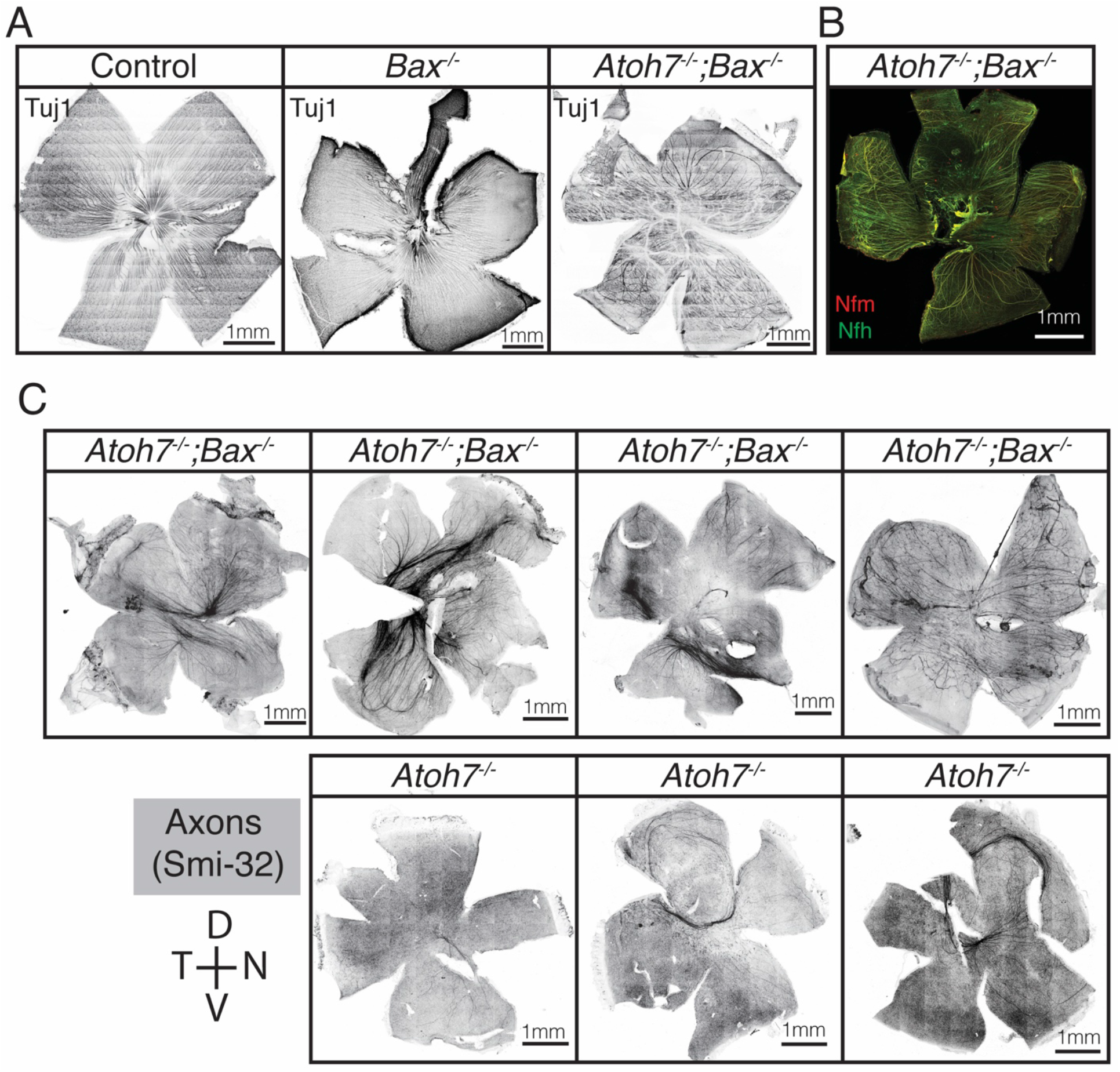
Peristimulus time histogram averaged from 120 repetition of 1-Hz square wave flash of every RGC recorded from in *Atoh7*^*−/−*^*;Bax*^*−/−*^ mice. The red line represents the transition from Light on to Light off in the square wave light flash.

**Supplemental Figure 5.**
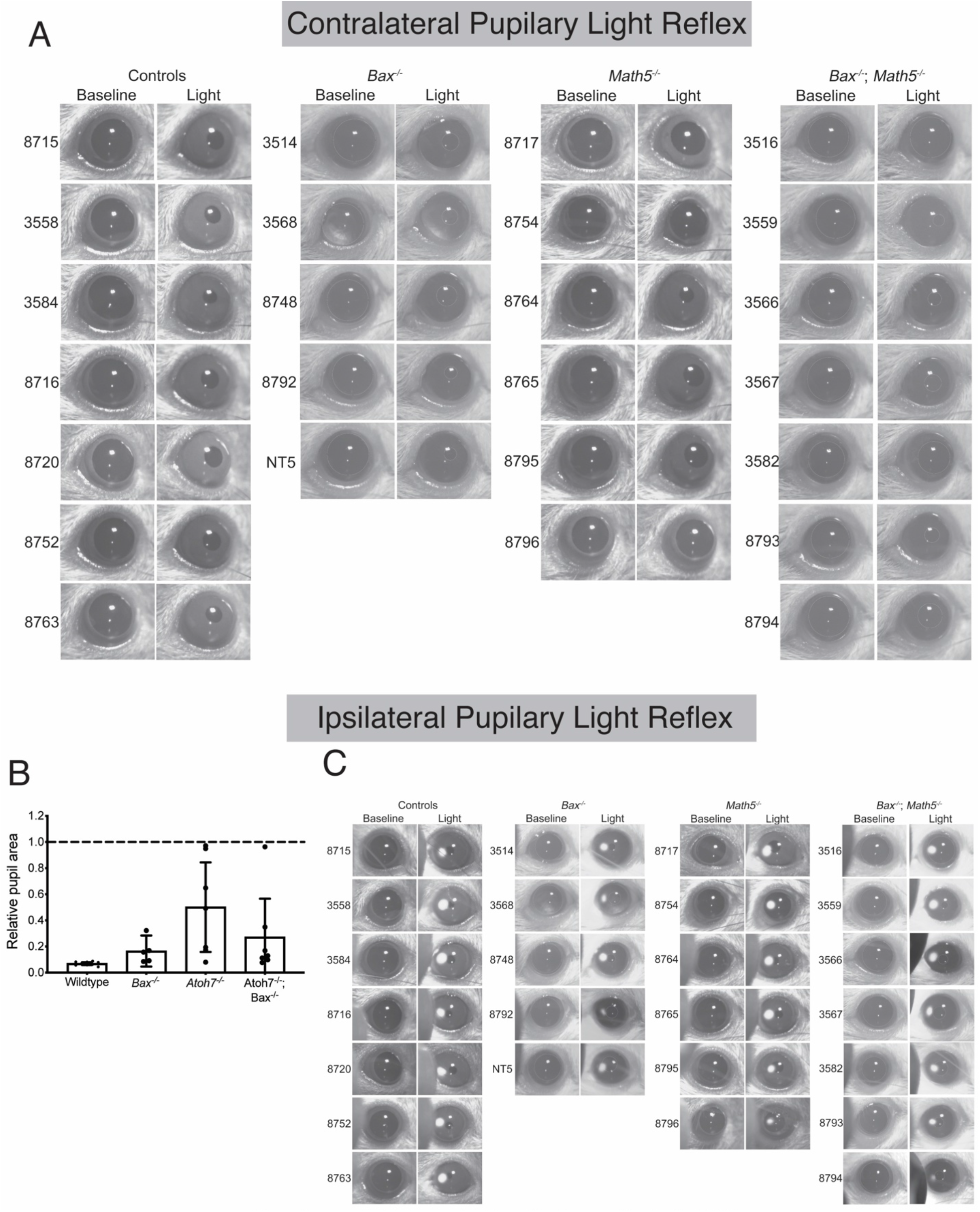
Similar results seen with Tuj1 (A), NFM, and NFH (B) immunostaining as seen with Smi-32 staining. (C) More examples of orientation preserved dissection of whole retinas of all four genotypes and stained with an antibody to Smi-32, notice the orientation of the retinas as marked by the orientation rose.

**Supplemental Figure 6.**
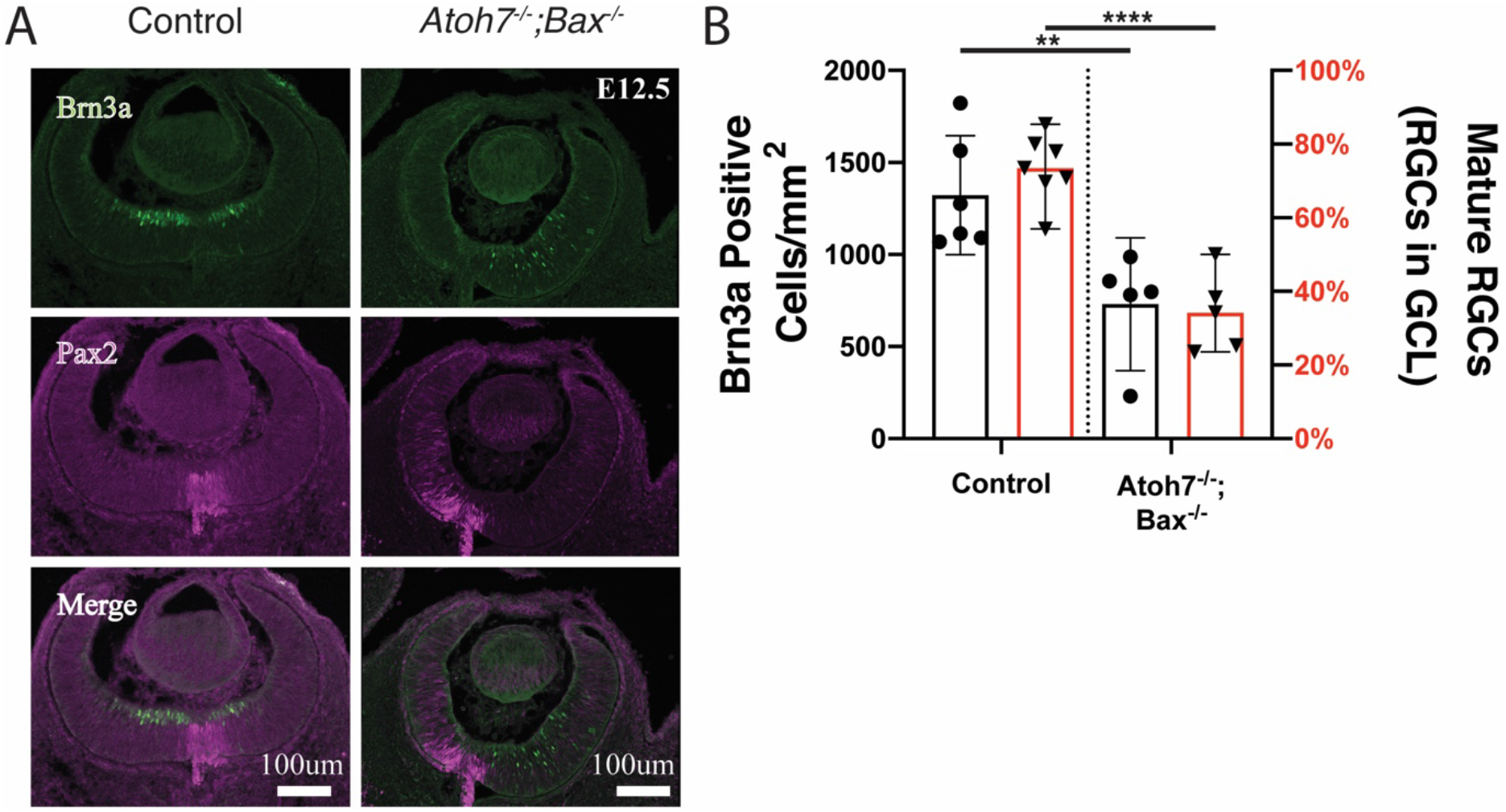
RGCs in *Atoh7*^*−/−*^*;Bax*^*−/−*^ mice do not sustain visual tasks. In the virtual optometer, animals visually track a moving grating of varied widths. (A) Only wild type animals were able to track gratings with an acuity of 0.4 cycles/degree. In the cued Morris Water Maze (B) animals locate a marked platform to escape the water. In four successive trials, most wild type animals learned to locate the platform relatively quickly. (C) ipRGC projections do not reach the brain in *Atoh7*^*−/−*^*;Bax*^*−/−*^ mice. The projections of ipRGCs were genetically labeled with the tauLacZ marker (depicted here as Opn4+/−) in the *Atoh7*^*−/−*^*;Bax*^*−/−*^ background. Robust innervation is revealed by X-Gal staining in the control while the *Atoh7*^*−/−*^*;Bax*^*−/−*^ mice do not show innervation (target nuclei outlined in red). (n=3, scale bars = 100mm). (D) Fluorescently conjugated Cholera toxin b labels RGC projections. Non-image forming (SCN, OPN) and image-forming targets (LGN, SC) do not receive retinal innervation in *Atoh7*^*−/−*^ or *Atoh7*^*−/−*^*;Bax*^*−/−*^ mice. Yellow outlines depict targets; All scale bars 100mm, n=3 per genotype. Mean ± 95% confidence intervals. Statistical significance tested by one-way ANOVA with Tukey’s post test for multiple comparisons * p=0.0386, **** p < 0.0001. Abbreviations: SCN: Suprachiasmatic nucleus, OPN: Olivary Pretectal Nucleus, LGN: Lateral geniculate nucleus, SC: Superior colliculus.

**Supplemental Figure 7.**
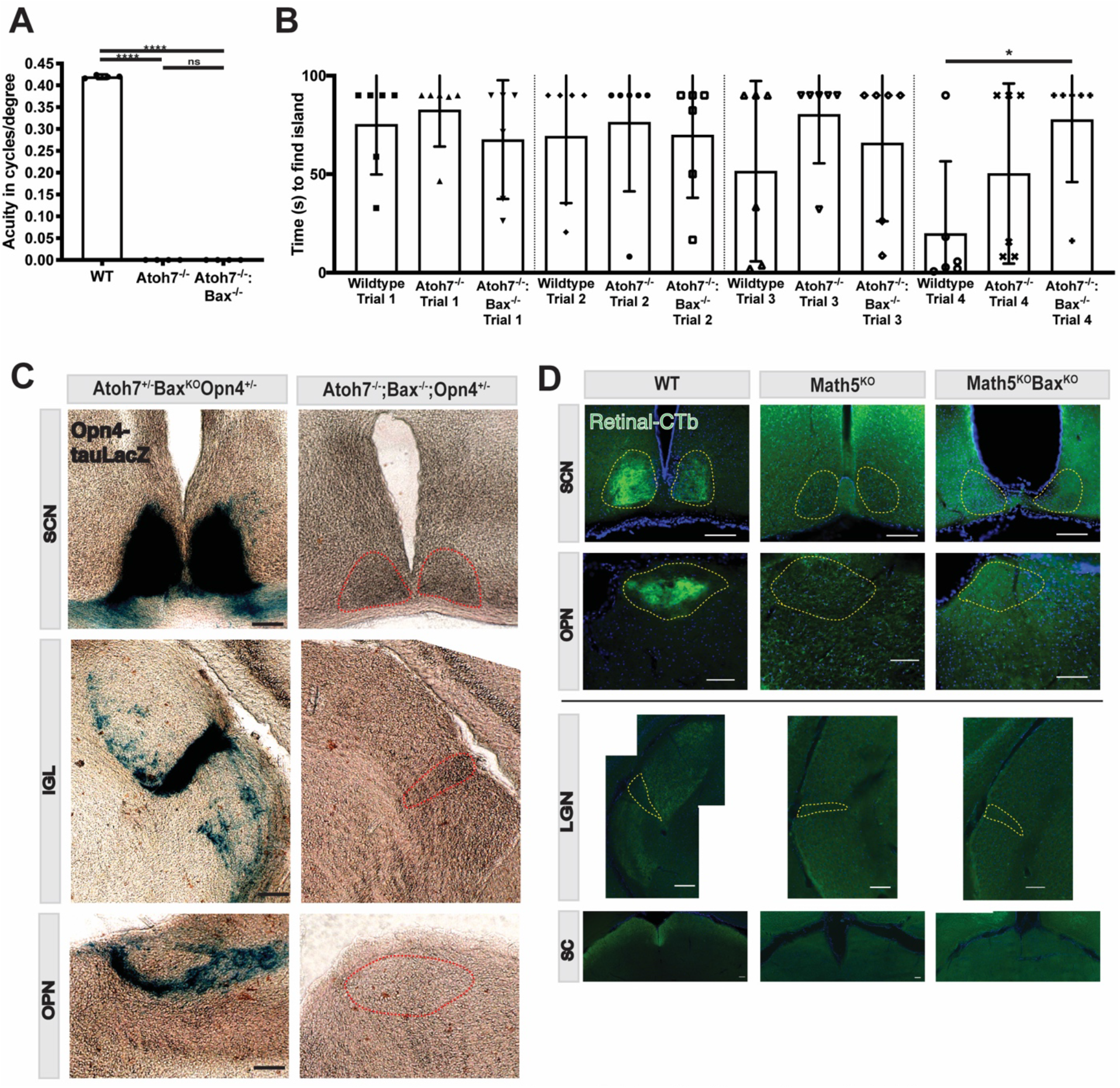
Traces of Pupillary light reflex (PLR) of individual mice before and after light stimulus for contralateral (A) and ipsilateral (D) recordings. *Bax*^*−/−*^ and *Atoh7*^*−/−*^*;Bax*^*−/−*^ mice lack retinal pigmentation therefore their pupils are outlined for clarity. Analysis of the relative pupil area of all of the PLR responses between genotypes for contralateral (B) and ipsilateral (C). Mean ± 95% confidence intervals.

**Supplemental Figure 8.**
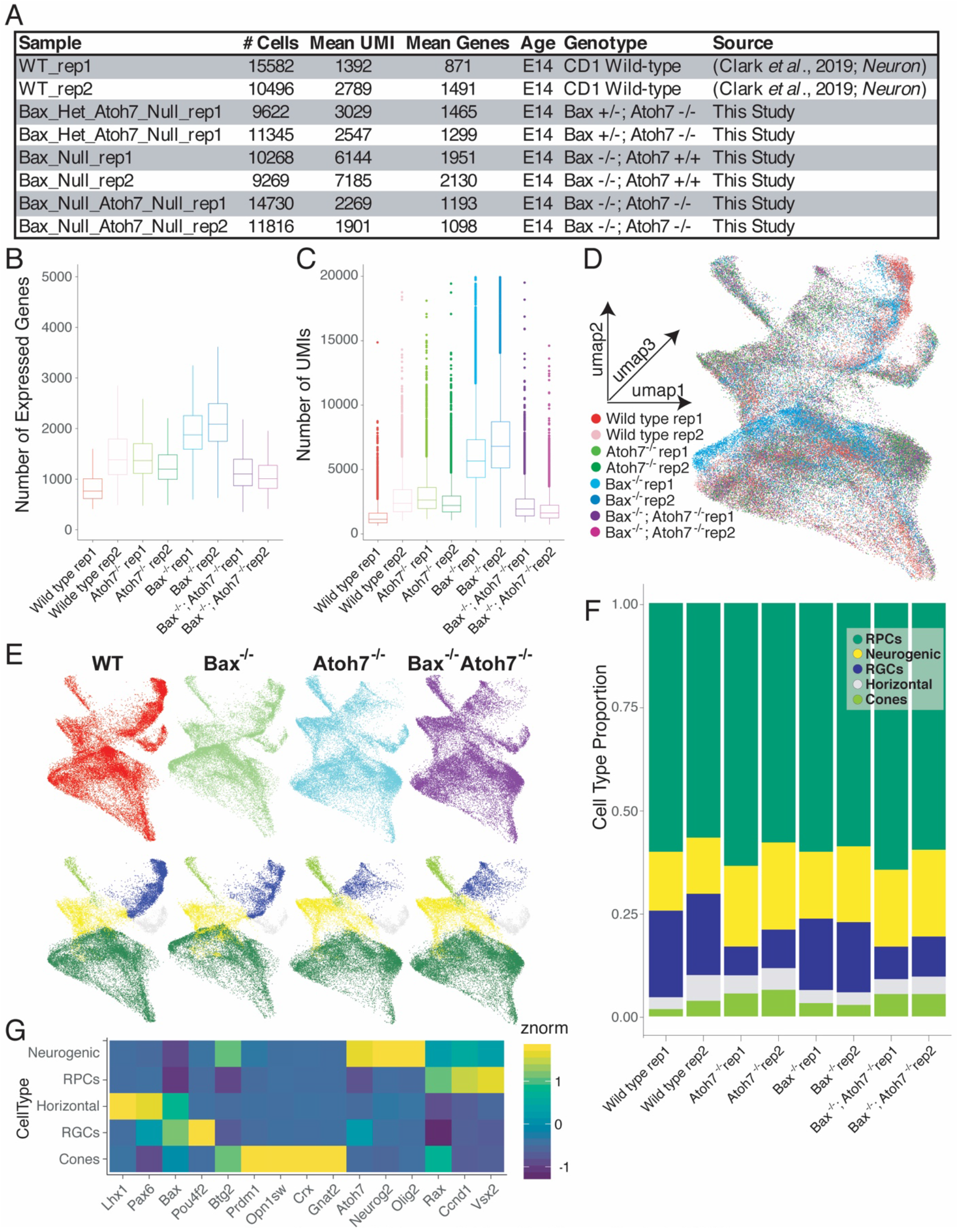
(A) Table of scRNA-seq statistics for each sample. (B-C) Boxplots of the number of (B) Expressed Genes and (C) Unique molecular identifier (UMI) or transcript counts within each cell. (D) UMAP dimension reduction of the aggregate single cell datasets with individual cells colored by originating sample identity. (E) UMAP dimension reductions displaying the subset of cells corresponding to each genotype (top) and the cell type annotations (bottom) of the corresponding cells. (F) Cell type annotations of cells within each scRNA-seq sample. (G) Heatmap of known cell type markers within annotated cell types.

**Supplemental Figure 9.**
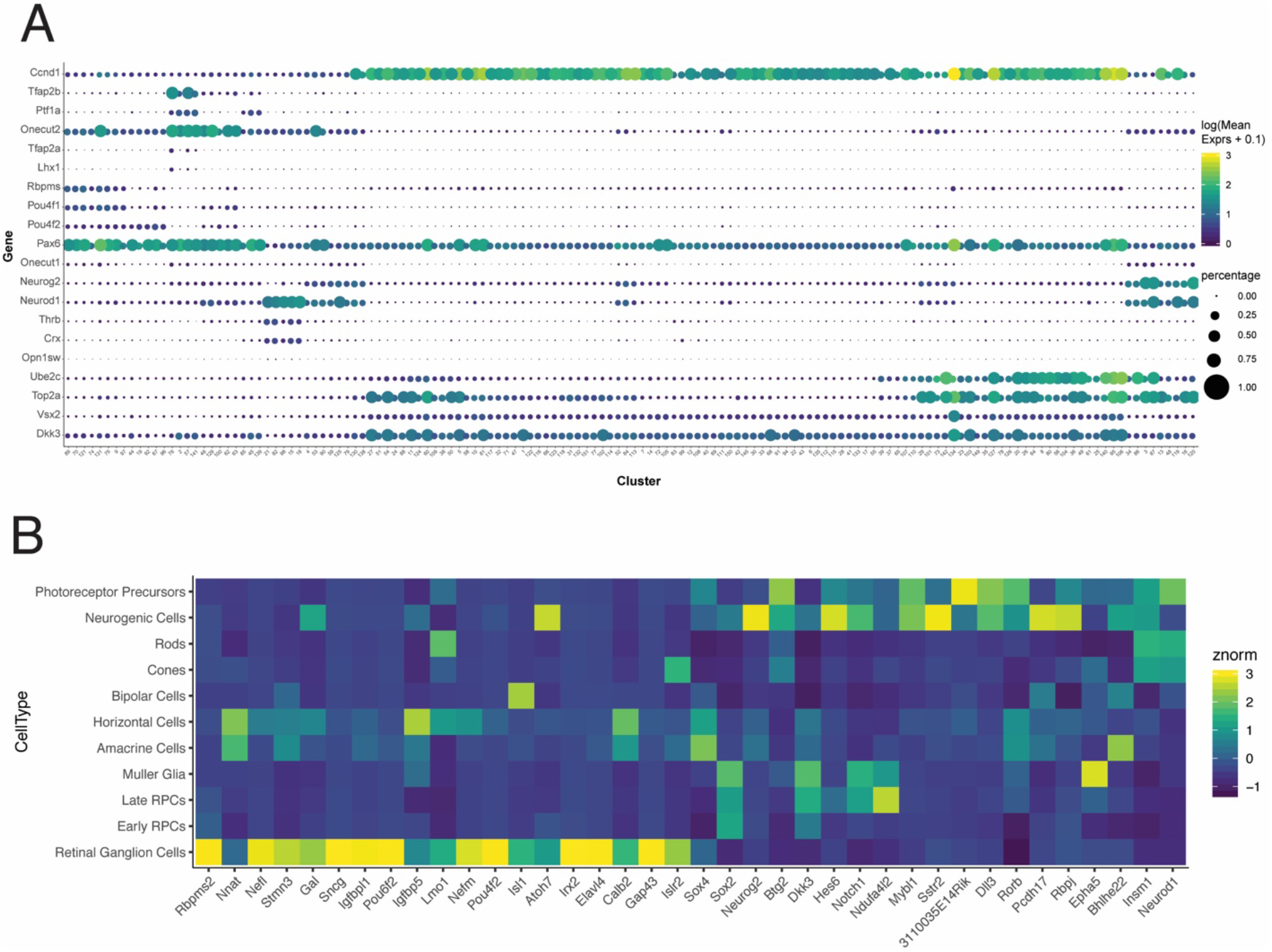
(A) Dotplot of a subset of marker genes used to determine cell type annotations of clusters within UMAP dimension reduction space. Color of the circle corresponds to the log() of the mean expression of individual transcripts within cells of each cluster. Size of the circles corresponds to the percentage of cells within the cluster that had at least one transcript detected. (B) Heatmap of the cell type enrichment of *Atoh7*-dependent, differentially expressed transcripts across wildtype mouse retinal development (E11-P14) as assayed by Clark et al., 2019.

**Supplemental Figure 10.**
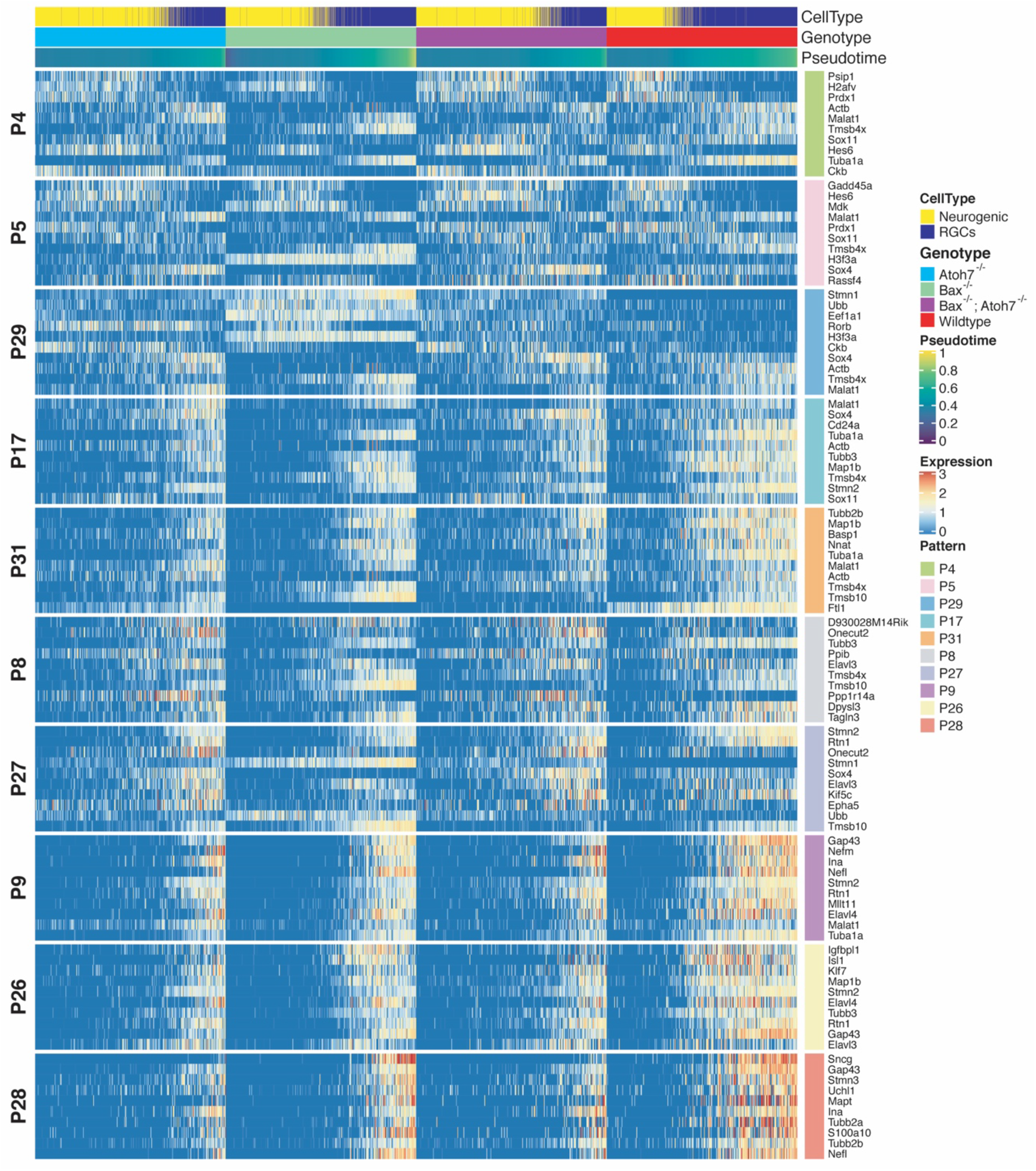
Pseudotime heatmap of top 10 weighted genes of scCoGAPS Patterns. Cells are ordered by pseudotime.

**Supplemental Figure 11.**
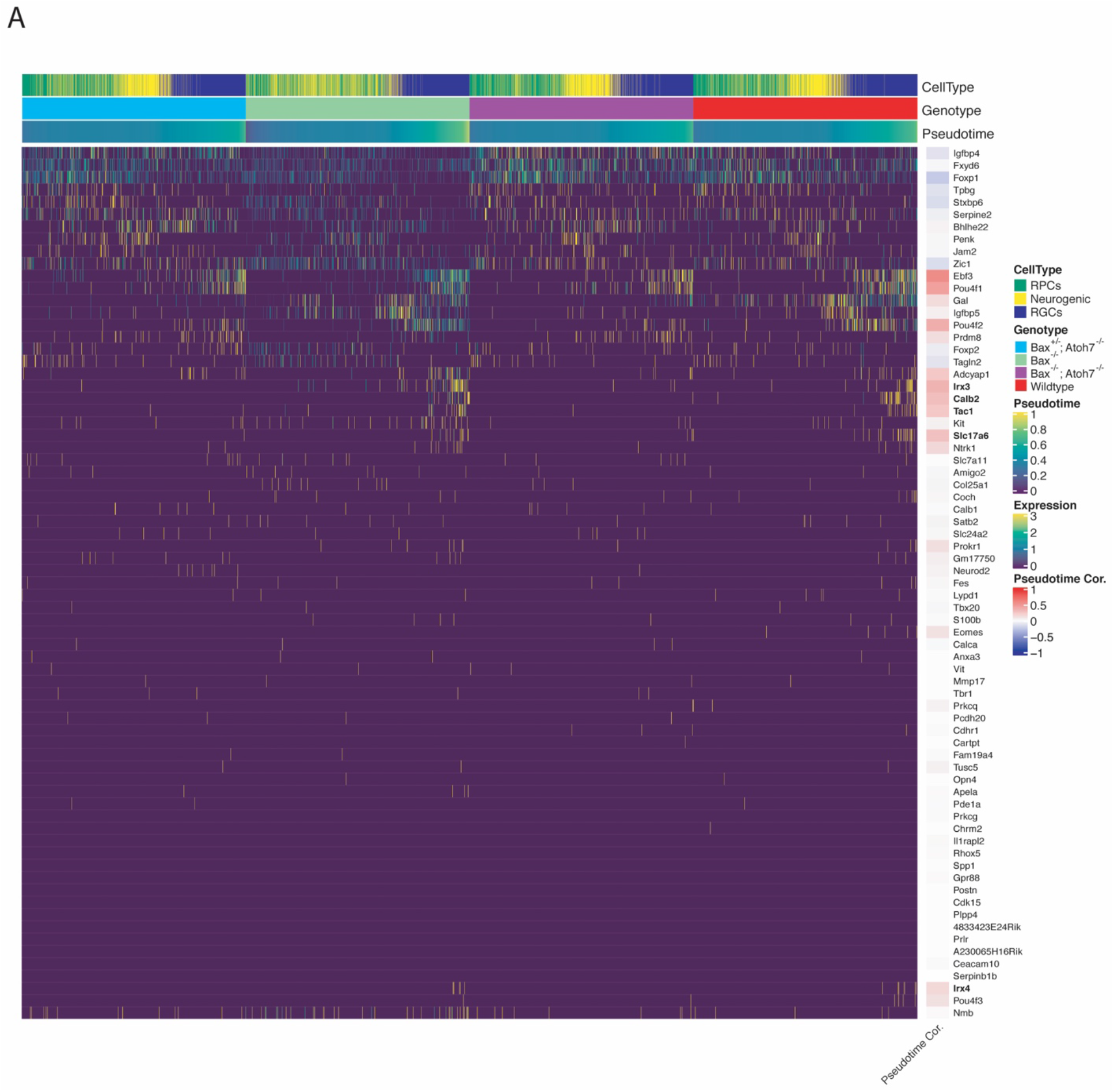
Pseudotime heatmap of previously published RGC subtype markers and their correlation to pseudotime.

**Supplemental Figure 12.**
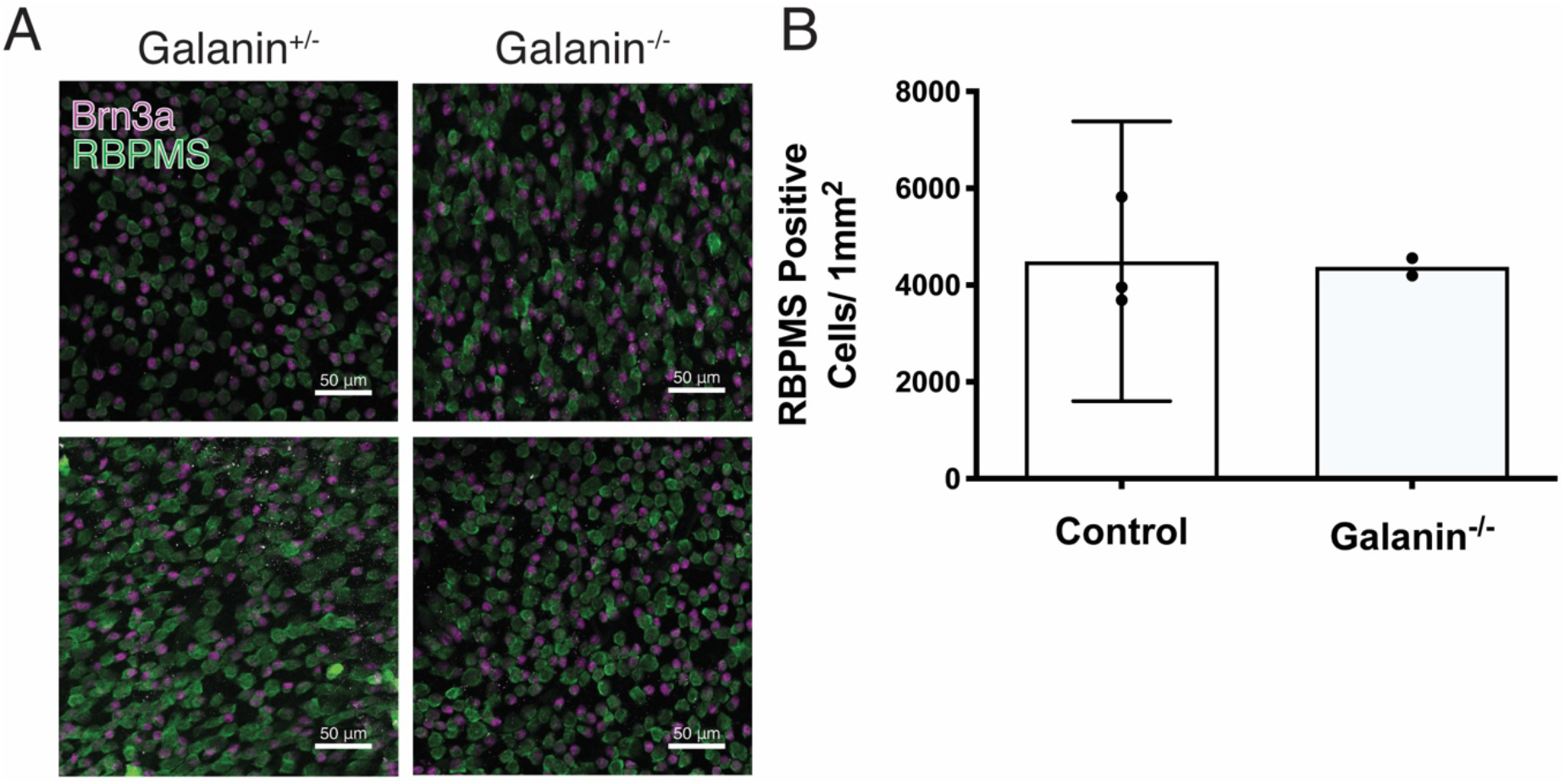
(A) Immunohistochemistry images of the RGC layer stained for Brn3a and RBPMS in Galanin heterozygous and knockout retinas. (B) Cell counts of RGCs indicate no difference in RGC number in P8 galanin-deficient retinas.

